# Distinct evolutionary trajectories of V1R clades across mouse species

**DOI:** 10.1101/2019.12.19.883082

**Authors:** Caitlin H Miller, Polly Campbell, Michael J Sheehan

## Abstract

**Background:** Many animals rely heavily on olfaction to navigate their environment. Among rodents, olfaction is crucial for a wide range of social behaviors. The vomeronasal olfactory system in particular plays an important role in mediating social communication, including the detection of pheromones and recognition signals. Currently, very few vomeronasal receptors have known ligands, which severely limits our understanding of chemosensory-driven social communication. In this study we examine patterns of vomeronasal type-1 receptor (V1R) evolution in the house mouse and related species within the genus *Mus*. By exploring the evolution of these receptors, we provide insight into the functional roles of receptor subtypes as well as the dynamics of gene family evolution.

**Results:** We generated transcriptomes from the vomeronasal organs of 5 *Mus* species, and produced high quality V1R repertoires for each species. We find that V1R clades in the house mouse and relatives exhibit distinct evolutionary trajectories. Some clades are highly conserved, while others reveal patterns of rapid evolutionary change. We identify putative species-specific gene expansions, including a dramatic clade D expansion in the house mouse. While gene gains are abundant, we detect very few gene losses. We describe a novel V1R clade and highlight candidate receptors for future de-orphanization. Based on clade-level evolutionary patterns, we identify receptor families that are strong candidates for detecting social signals and predator cues. Our results further support the view that V1Rs are important for detecting the physiological status of conspecifics, particularly female estrus cues.

**Conclusion:** Analysis of clade-level evolution is critical for understanding species’ chemosensory adaptations. This study provides clear evidence that V1R clades are characterized by distinct evolutionary trajectories. As receptor evolution is shaped by ligand identity, these results provide a framework for examining the functional roles of different receptors.

## BACKGROUND

Olfaction involves detecting and discriminating among chemicals in the environment. Chemical compounds can vary considerably in structure, creating a highly complex chemical space in which olfactory systems evolve. In most mammals, olfaction relies on two discrete receptor systems: the main olfactory receptors (ORs) and the vomeronasal receptors (VRs) (Dulac & Torello, 2003; Meisami & Bhatnagar, 1998; Restrepo et al., 2004). ORs detect a broad range of environmental odors (Liberles, 2014; Mombaerts, 2004; Nara et al., 2011), while VRs are integral to species-specific chemical detection, including pheromone detection (Boschat et al., 2002; Dulac & Axel, 1995). In humans, ORs are the only olfactory receptors, as the vomeronasal system is no longer functional. In other species, however, VRs mediate a wide range of social behaviors, including sexual, aggressive, and parental behaviors (Chamero et al., 2007; Ferrero et al., 2013; He et al., 2008; Kaur et al., 2014; Kimoto et al., 2005; Leypold et al., 2002; Orikasa et al., 2017; Powers & Winans, 1975; Stowers et al., 2002; Tachikawa et al., 2013). VRs thus provide a unique window into the chemical basis of social behaviors and the evolution of pheromone detection.

Across species, VRs exhibit striking evolutionary patterns. Whereas ORs have largely orthologous relationships among divergent species (Grus & Zhang, 2008), VR evolution is characterized by rapid gene turnover wherein receptors are quickly gained and lost over evolutionary time (Grus & Zhang, 2004; Jiao et al., 2019; Lane et al., 2004). This pattern of gene birth-and-death results in lineage-specific receptor repertoires (Grus & Zhang, 2008). Consequently, there are substantial differences in receptor sequences and repertoire size across divergent species (Grus et al., 2005; Jiao et al., 2019; Shi et al., 2005; Silva & Antunes, 2017; Yang et al., 2005; Young et al., 2010). For example, among three mammalian species (dog, opossum, and house mouse) there are virtually no one-to-one VR orthologs (Grus & Zhang, 2008). This is perhaps not surprising given the broad evolutionary timescale examined. However, even among two rodent species (the rat and house mouse), the majority of VRs fall into lineage-specific clades with very few orthologs detected (Grus & Zhang, 2004).

As one of the leading model organisms, further understanding chemosensation in the house mouse will provide insight into how chemosensory stimuli mediate distinct behavioral and neural responses. House mice are valuable models for examining VRs as they have large VR repertoires and there exists a wealth of knowledge on their social behavior, neural activity, and genetics (Brignall & Cloutier, 2015; Chamero et al., 2007; Dulac & Wagner, 2006; Duyck et al., 2017; Ferrero et al., 2013; Ibarra-Soria et al., 2014; Kaur et al., 2014; Orikasa et al., 2017; Tachikawa et al., 2013; Wagner et al., 2006; Wynn et al., 2012). Currently, very few VRs have known ligands, which presents a significant barrier to studying the mechanisms underlying social behavior in house mice (Haga-Yamanaka et al., 2014; Haga et al., 2010). By examining the evolutionary trajectories of VRs, we may uncover distinct evolutionary patterns among receptor clades, and thereby identify targets for study based on the extent of turnover or conservation observed in a clade.

Vomeronasal sensory neurons express two major gene families in a cell-specific manner: V1Rs (type-1 VRs) and V2Rs (type-2 VRs) (Mombaerts, 2004; Silva & Antunes, 2017). V1Rs consist primarily of single-exon genes whereas V2Rs are multi-exonic (Rodriguez, 2016). Structurally, V1Rs have a short N-terminal extracellular region whereas V2Rs have long and highly variable N-terminal domains (Mombaerts, 2004; Rodriguez, 2016). We focused on V1Rs in this study due to the genetic tractability of their simpler gene structure for transcriptome assembly and sequence analysis. In functional terms, V1Rs primarily detect airborne volatiles (Chamero et al., 2007; Del Punta et al., 2002; Kimoto et al., 2005; Leinders-Zufall et al., 2000; Leinders-zufall et al., 2004; Meeks et al., 2010). In house mice, V1Rs have been implicated in detecting a wide range of volatiles, including urinary steroid molecules that are crucial for gender discrimination and sexual behaviors (Haga-Yamanaka et al., 2014; Isogai et al., 2011; Nodari et al., 2008; Novotny et al., 1986; Novotny et al., 1999).

Here, we characterize patterns of V1R evolution among the house mouse and relatives. We take a molecular evolutionary approach and analyze V1R repertoires across six species within the genus *Mus* (**Figure 1**): *M. m. domesticus* (house mouse), *M. spicilegus, M. macedonicus, M. spretus, M. caroli*, and *M. pahari*. By examining the under-explored timescale of VR evolution among closely related species, this dataset offers new insight into the dynamics of VR evolution and provides a framework for understanding the selective pressures shaping V1R clades. Examining the evolutionary history of V1R clades may in turn guide future efforts to deorphanize receptors in the house mouse. For example, receptors that detect crucial chemical cues, such as predator odors or sex-specific signals, are likely conserved across *Mus* species. Conversely, receptors involved in detecting variable signals within or between species, such as identity signals or pheromones, may display species-specific evolutionary patterns. The house mouse is the only *Mus* species commensal with humans, and may exhibit gene gains or losses that reflect chemosensory adaptations toward their unique commensal ecology (Pocock et al., 2004; Suzuki et al., 2013). Thus, different V1R genes may experience distinct selective pressures that reflect ligand function. Examining evolutionary patterns of V1Rs can guide functional studies of receptors, as well as provide general insights into sensory evolution.

**Figure 1.**
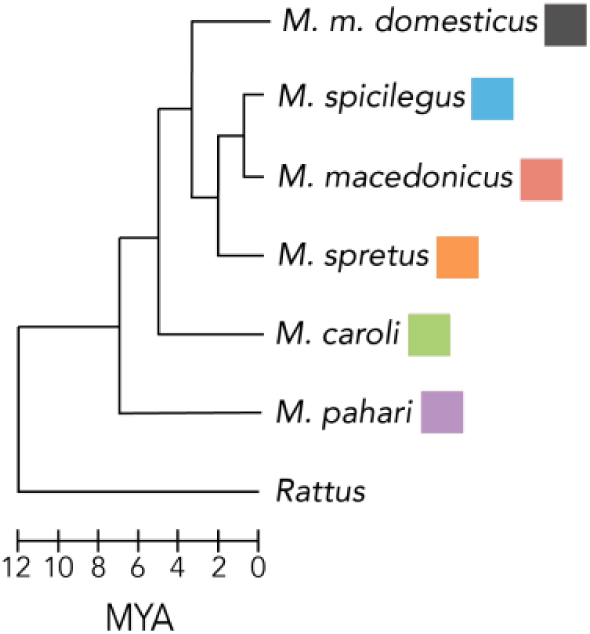
*Mus* species phylogeny. Includes all species used in this study. Rat (*Rattus norvegicus*) provided as an outgroup. Modified from Steppan & Schenk (2017). Divergence estimates from Chevret et. al. (2005). Species colors used throughout.

## RESULTS

### VNO Sequencing, Assembly & V1R Recovery

We characterize V1R repertoires for five *Mus* species of varying evolutionary distance from the house mouse (1.5-7 mya, **Figure 1**) by sequencing their VNO transcriptomes using short-read platforms. The final transcriptome assemblies for each species are of good quality (**Table 1**). We detect approximately twice the number of V1Rs than are currently annotated in the genomes of *M. spretus, M. caroli*, and *M. pahari* and provide the first *M. macedonicus* V1R dataset (**Table 1**). The number of V1Rs identified in *M. spicilegus* is in good agreement with existing genome annotations (**Table 1**). For one species (*M. spretus*), the short-read sequencing was performed at greater depth, and an additional round of long-read sequencing was done. This allows us to examine the effectiveness of short versus long-read sequencing for assembling large and highly duplicated gene families such as V1Rs. The total number of assembled transcripts is greater for the *M. spretus* short-read dataset, as expected from greater sequencing depth (**Table 1**).

**Table 1.** VNO transcriptome assembly statistics, V1R transcript recovery, and genome annotations. The mouse reference genome is shown for comparison (*M. m. domesticus*, top). Recovery estimates combining short and long-read datasets for *M. spretus* are indicated in bold.

| Species | Total Transcripts | Mean Length (bp) | N50 | % with ORF<sup>f</sup> | Total V1R Transcripts | V1Rs With Unique Annotations | V1R Genes Detected (With Duplicates) | Genome Annotated V1Rs |
| --- | --- | --- | --- | --- | --- | --- | --- | --- |
| *M. m. domesticus*<sup>a</sup> | --- | --- | --- | --- | 263 | --- | 208 | 208 |
| *M. spicilegus*<sup>b</sup> | 228,809 | 664 | 1632 | 43 | 122 | 105 | 119 | 120 |
| *M. macedonicus* | 255,395 | 649 | 1630 | 42 | 129 | 117 | 126 | --- |
| *M. spretus*<sup>c</sup> | 1,181,673 | 688 | 1587 | 31 | 134 (253) | 108 (146) | 180 | 85 |
| *M. caroli*<sup>d</sup> | 384,865 | 460 | 1919 | 42 | 131 | 110 | 126 | 50 |
| *M. pahari*<sup>e</sup> | 450,181 | 305 | 3107 | 45 | 117 | 93 | 113 | 45 |

<sup>a</sup>GRCm38.p6, <sup>b</sup>GenBank accession#: QGOO00000000, <sup>c</sup>SPRET\_EIJ\_v1.1, <sup>d</sup>CAROLI\_EIJ\_v1.1, <sup>e</sup>PAHARI\_EIJ\_v1.1, <sup>f</sup>ORF: open reading frame

On average, 126 V1R transcripts are recovered from each species’ short-read assembly (**Table 1**). A subset are transcript variants or gene duplicates, with homology to the same gene in the mouse reference genome (GRCm38.p6). Although the majority of V1Rs are single-exon genes, a substantial number of V1Rs contain introns and express transcript variants in the house mouse (Ibarra-Soria et al., 2014), we similarly detect transcript variants among the non-commensal species sequenced (**Table 1 & Figure 2**). For a conservative estimate of V1R genes, only unique transcript annotations are included (**Table 1**). When putative gene duplicates are added, the number of V1R genes increases markedly (**Table 1**). Compared to the house mouse, the 5 *Mus* species sequenced have smaller V1R repertoires, consistent with V1R gene expansion in the house mouse (**Table 1**). However, the addition of long-read sequencing for *M. spretus* increases the number of V1Rs genes detected, resulting in a repertoire size similar to the house mouse (**Table 1**). Therefore, whereas the *M. spretus* V1R repertoire is likely close to complete, long-read sequencing may detect additional V1Rs in *M. spicilegus, M. macedonicus, M. caroli* and *M. pahari*. Importantly, our analysis of V1R evolution in *Mus* is based on (1) a well-annotated mouse reference genome, (2) a comprehensive *M. spretus* V1R dataset, and (3) >100 V1Rs for all 6 *Mus* species. Therefore, small gaps in detection across the entire V1R family should not bias the broad patterns of V1R evolution reported here. Furthermore, discrepancies in species’ repertoire size appear to be largely accounted for by a house mouse specific V1R gene expansion, discussed in further detail below.

**Figure 2.**
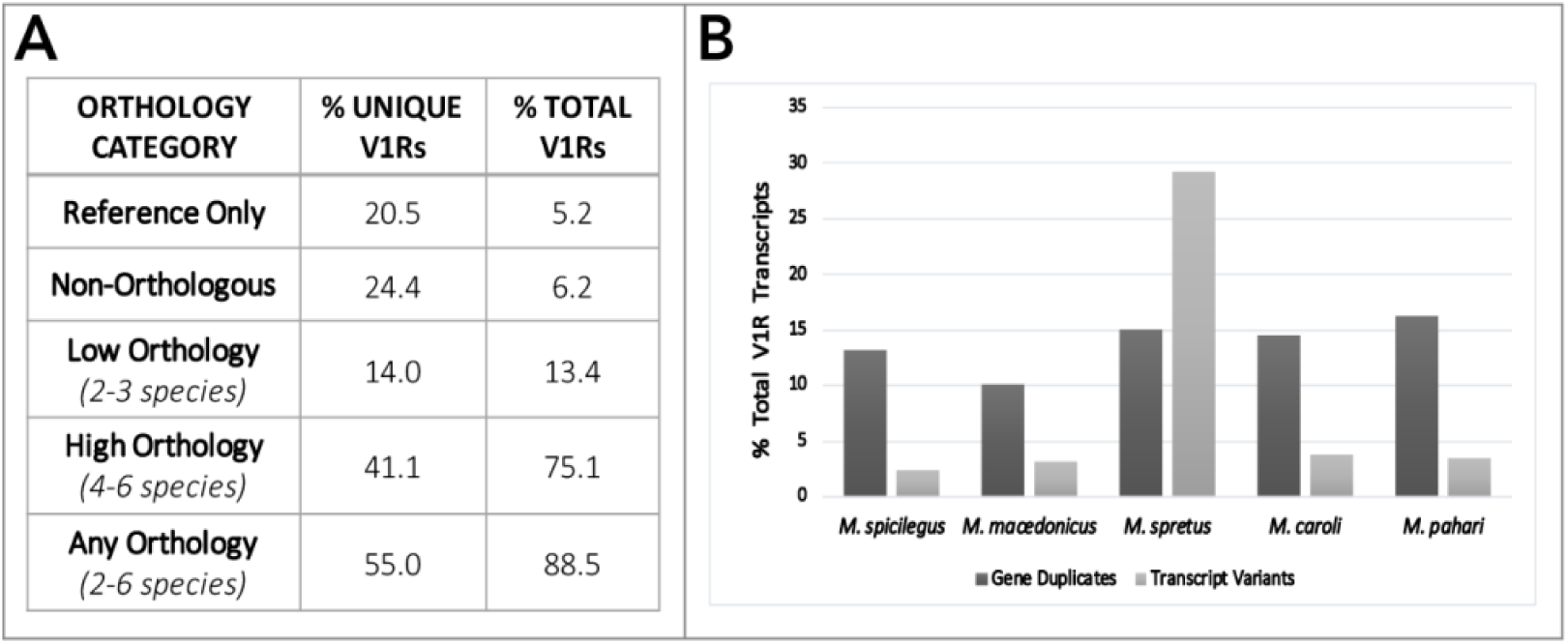
V1R orthology, gene duplicates, and transcript variants across *Mus* species. **(A)** The percent of *unique* and *total* V1Rs in each orthology category. **(B)** The percent of V1R transcripts that are either putative gene duplicates or transcript variants, for each species sequenced.

### V1R Evolution Across *Mus* Species

To explore V1R evolution, we characterize which receptors share a common ancestor (i.e. are orthologous) by examining relationships within a V1R gene tree containing all six *Mus* species (5 sequenced species and the house mouse reference (**Additional File 1**). A subset of receptors for each non-commensal mouse species do not exhibit a clear orthologous relationship to any V1R annotated in the mouse reference genome and are classified as non-orthologous genes (**Figure 2**). Similarly, a set of receptors annotated in the mouse reference genome was not detected in any other species (**Figure 2**).

We classify V1Rs into three broad categories based on their orthologous relationships: (1) V1Rs present only in the mouse reference genome, (2) non-orthologous V1Rs, and (3) V1Rs with orthology across multiple species. V1Rs with orthology across multiple species are further categorized based on the number of species represented in each orthologous receptor group (orthogroup). Orthogroups with 2-3 species were classified as “low orthology,” and orthogroups with 4-6 species as “high orthology” (**Figure 2A**). The majority of transcripts have some evidence for orthology (88.5%, **Figure 2A**). Furthermore, most transcripts are highly orthologous (74.8%, **Figure 2A**), indicating that missing V1Rs are unlikely to bias broad patterns identified here. Although many receptors are shared across species, approximately 25% of all V1R transcripts, and 59% of all unique V1R annotations, are either low orthology, non-orthologous, or present only in the mouse reference genome (**Figure 2A**). This indicates that the dramatic VR gene turnover observed among more divergent mammalian species, such as across tetrapods or between rodent species (Grus & Zhang 2004; Grus & Zhang, 2008; Silva and Antunes, 2017), is replicated within the genus *Mus*, albeit on a more limited scale.

We next examine the presence of gene duplicates and transcriptional variation across species (**Additional File 2**). A similar proportion of V1R gene duplicates are identified across all 5 species (10-16%, **Figure 2B**). The proportion of V1R transcript variants detected is also similar across species, with the clear exception of *M. spretus* (**Figure 2B**). As expected, the addition of long-read (*M. spretus*) sequencing data recovers many more transcript variants than short-read sequencing datasets (**Figure 2B**). Interestingly, the same number of V1R genes expressing distinct coding transcript variants are detected in *M. spretus* as in the house mouse (43 V1R genes, **Additional File 3: Figure S1**). However, the identity of V1Rs exhibiting alternative spliceforms, and the clades they belong to, vary between the two species (**Additional File 3: Figure S1**). In contrast, the proportion of gene duplicates detected is similar between *M. spretus* and the other species. This indicates that, for gene families such as V1Rs, short-read datasets are sufficient for identifying gene duplicates.

Our characterization of V1R repertoires across *Mus* species allows for a reliable estimate of V1R gene loss in the house mouse. We detect evidence for 10 such V1R gene losses, distributed across six clades (**Table 2** & **Figure 3A**: indicated in red text). All V1R genes lost in the house mouse are present in at least 3 of the 5 non-commensal *Mus* species examined, including close relatives (**Table 2**). Most gene losses have corresponding pseudogenes in the house mouse reference genome (**Table 2**). It appears gene losses are relatively uncommon compared to the abundant gene gains, at least within the house mouse lineage.

**Table 2.**
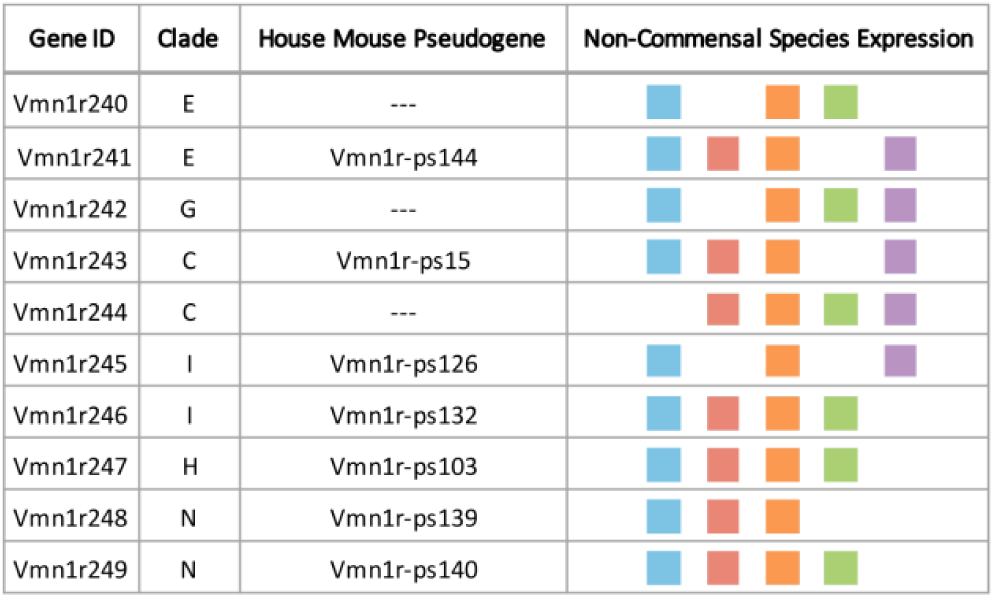
V1R gene losses in the house mouse. Species with expression indicated with different colors: 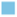 *M. spicilegus*, 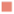 *M. macedonicus*, 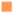 *M. spretus*, 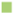 *M. caroli* and, 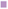 *M. pahari*.

**Figure 3.**
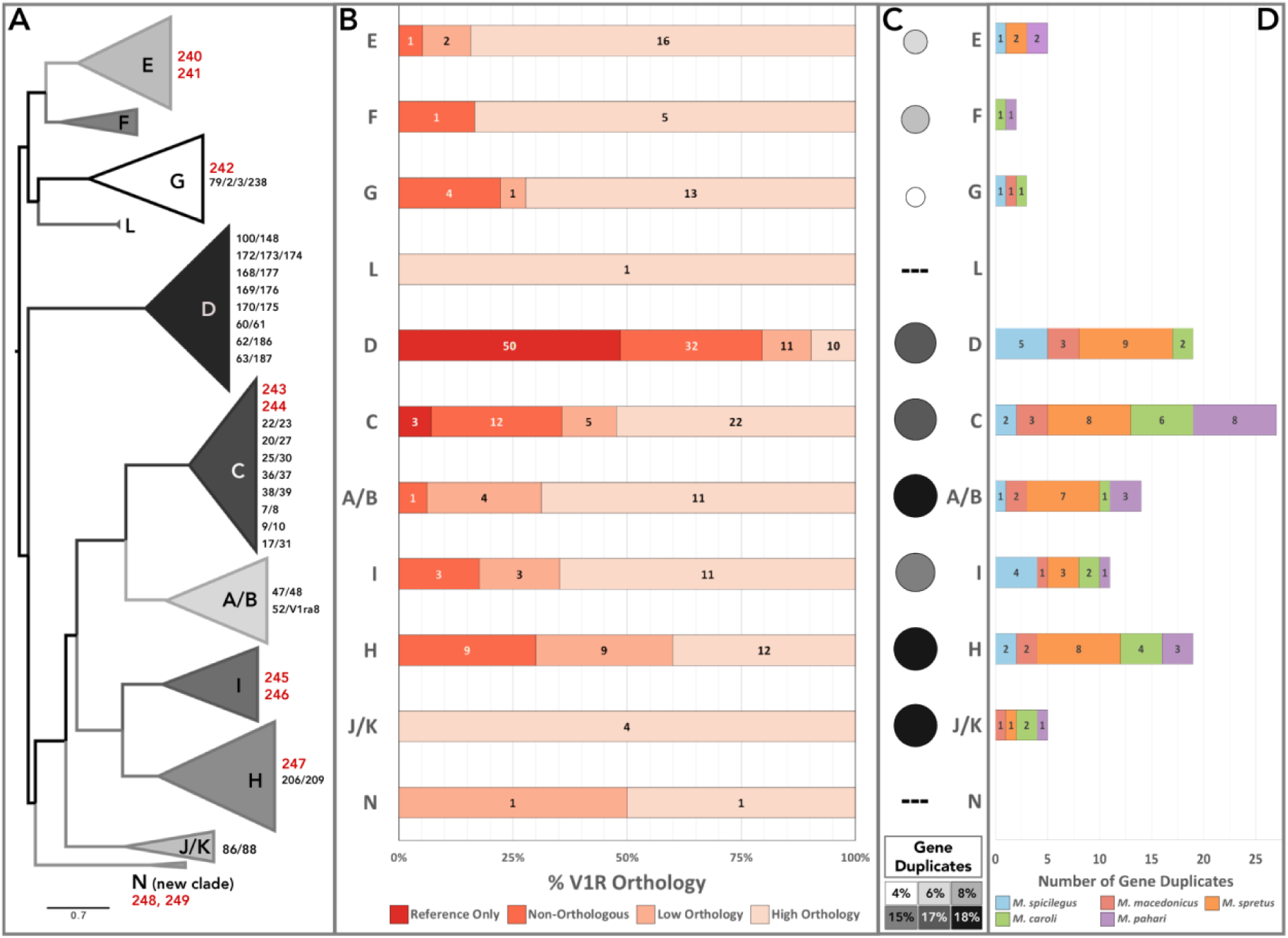
Patterns of orthology and gene duplication across V1R clades. **(A)** Phylogeny of all 11 V1R clades. Scale bar indicates 0.7 amino acid substitution per site. Shown to the right of each clade: V1R gene losses in the house mouse (red), and orthogroups with multiple reference annotations (combination-IDs, black). **(B)** Percent of V1Rs by clade that fall into each orthology category: present only in the reference-only, non-orthologous, low orthology and high orthology. **(C)** Percent of sequenced transcripts (reference not included) that are gene duplicates. **(D)** Total number of V1R gene duplicates detected in each clade for each species.

### Novel V1R Clade: Clade “N”

In addition to the house mouse gene losses observed in clades E, C, H, I and G, we identify a novel V1R clade (**Table 2, Figure 3A**). This novel clade “N” has been lost in the house mouse and consists of two receptor orthogroups. Both clade N receptors (*Vmn1r248* and *Vmn1r249*) are expressed in at least 3 non-commensal *Mus* species (**Additional File 3: Figure S2**) and have corresponding pseudogenes in both the house mouse (*M. m. domesticus*) and the rat (*Rattus norvegicus*).

### Variable Patterns of Evolution Among V1R Clades

Patterns of V1R gene orthology and duplication vary across clades. Four of the 11 V1R clades are highly orthologous (E, F, J/K and L: >75% of receptors are high-orthology), with clade G trailing just behind with a few more non-orthologous receptors (**Figure 3A, B**). All 5 of these clades have 5 or fewer gene duplicates detected, however, the proportion of duplicates by clade size is variable (**Figure 3C, D**). Clades E, F and G have very low proportions of gene duplicates, while clade J/K has among the highest (**Figure 3C**).

Clades C, D and H have abundant low-orthology and non-orthologous receptors (**Figure 3B**), indicating greater evolutionary lability. While most orthologous relationships are straightforward, some orthogroups contain multiple house mouse receptors, and are annotated with combination-IDs (e.g. *Vmn1r25/30*). These receptor groups are the result of one or more duplication events within the *Mus* lineage, and are unequally distributed across clades, with 76% located in clades C and D (**Figure 3A**). In addition, all reference-only V1Rs are located in these same two clades (**Figure 3B**). Not surprisingly, clades C, D and H have the highest number of detected gene duplicates (19 or more) and have similarly high proportions of duplicates by clade size (**Figure 3C, D**). Thus, all three clades have evidence for substantial gene expansions, particularly clade D within the house mouse lineage.

We examine V1R clade sizes across all 6 species. With the striking exception of clade D, the house mouse clade sizes are very similar to the 5 other species, (**Figure 4**). This general pattern provides further evidence that receptor recovery is high and species’ repertoires are near complete. Interestingly, the *M. spretus* repertoire is largest for several clades (A/B, C, E, H and I; **Figure 4**), indicative of *M. spretus*-specific gene expansions.

**Figure 4.**
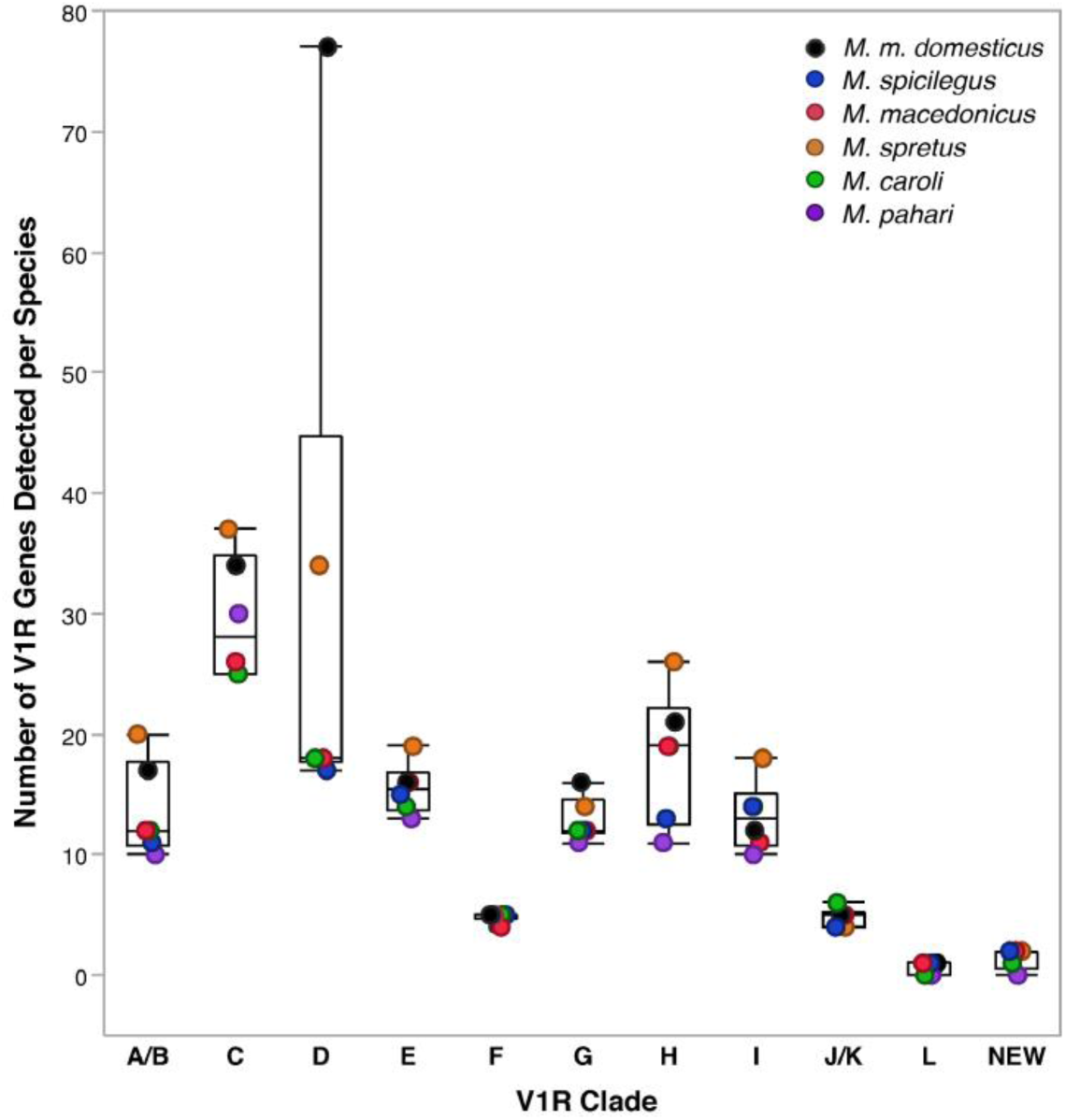
V1R clade receptor repertoire sizes across *Mus* species. *Mus* species are indicated with different colors.

The size ranges of two clades (A/B and D) are skewed by the house mouse and *M. spretus* datasets. Both species have much larger clade D repertoires than the other 4 species, exposing this clade as a potential hotspot for recent gene duplications (**Figure 4**). While the current data suggest these expansions are unique to *M. spretus* and *M. domesticus*, additional long-read sequencing might reveal comparable patterns in the other species. In contrast, there are several clades which exhibit low variation in repertoire size across all species’ datasets (E, F, G, J/K, L, N). Furthermore, clades C and H display variation in repertoire size across all 6 species, providing evidence for species-specific V1R gains and losses in multiple *Mus* lineages (**Figure 4**).

Guided by the evolutionary patterns observed across clades, we identify and categorize receptors as interesting candidates for further functional work based on striking patterns of conservation or divergence (**Additional File 3: Table S1**). We hope this list will help guide future efforts to deorphanize V1Rs.

### Fast-Evolving Clades & Lineage-Specific Expansions

#### Clade H

Clade H appears to be a mouse-specific V1R expansion, as it’s absent in the rat genome (Grus & Zhang, 2004). The clade is characterized by patterns of low orthology, abundant gene duplicates, and variable repertoire size across species (**Figures 3 & 4**). A sub-region of clade H containing *Vmn1r217, 219* and *220* receptors exemplifies the pattern of low orthology, while the receptor ortholog group *Vmn1r206/209* is representative of the abundant gene duplicates (**Figure 5A**). A striking exception to the patterns of dynamism observed in clade H is the extremely conserved receptor group *Vmn1r197* (**Figure 5A**). The general pattern of low orthology and the rapid species-specific gene gains and losses, suggests that clade H receptors may play an important role in species-specific chemosensation.

**Figure 5.**
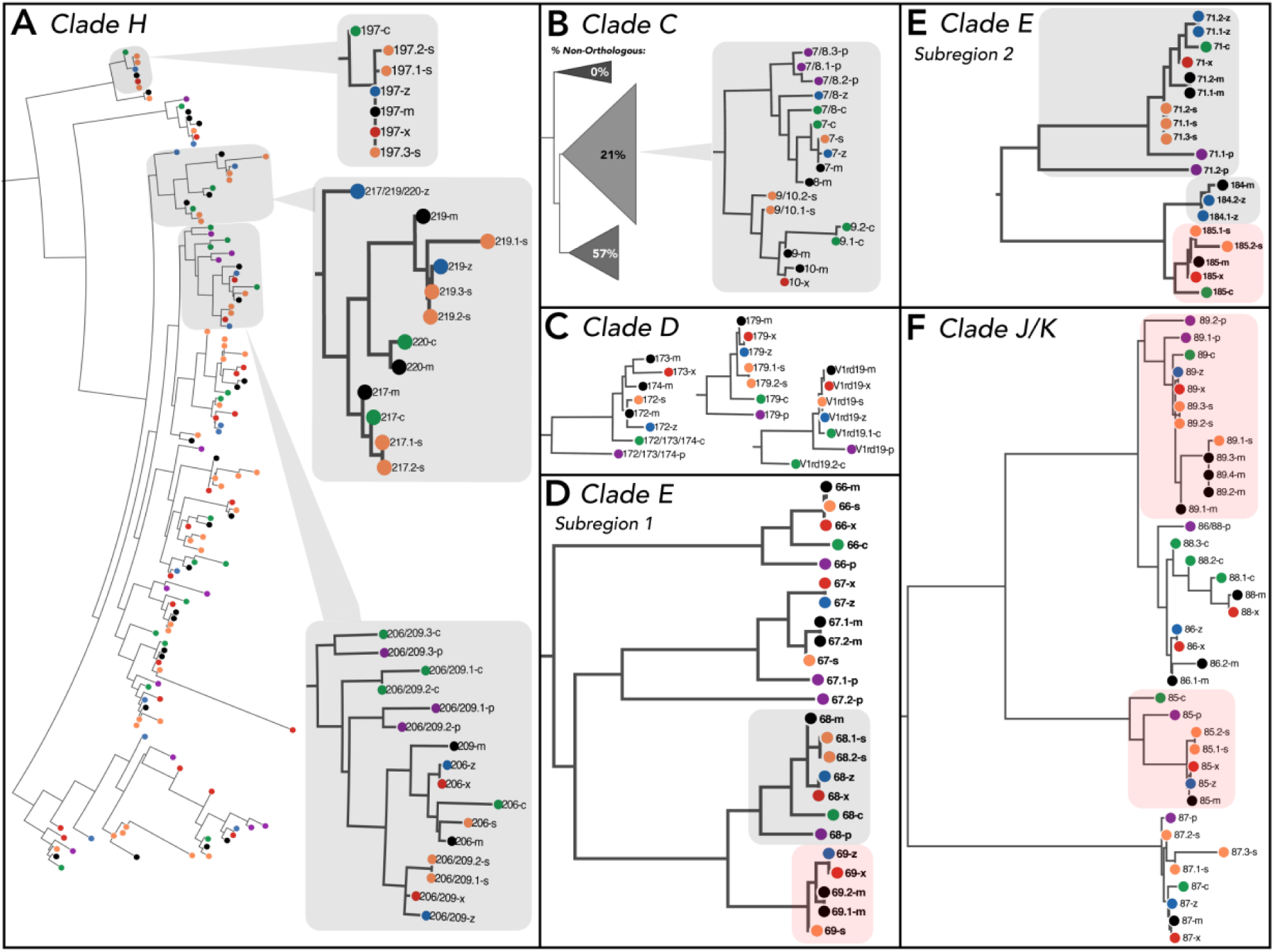
Example receptor groups depicting patterns of lineage-specific evolution and conservation across clades. V1R annotations are abbreviated (e.g. *Vmn1r137* as 137). Species transcripts annotated with the same reference gene have unique transcript IDs (e.g. 217.1 and 217.2). *Mus* species are indicated with colors and letters (*M. m. domesticus*: “m” and black; *M. spicilegus*: “z” and blue; *M. macedonics*: “x” and red; *M. spretus*: “s” and orange; *M. caroli*: “c” and green; *M. pahari*: “p” and purple). **(A) Clade H**: entire clade depicted, highlighting receptor groups that depict patterns of conservation and divergence. **(B) Clade C:** sub-clades shown with percentages of non-orthologous receptors. *Vmn1r7/8* and *Vmn1r9/10* are shown in detail. **(C) Clade D:** specific clade D receptor groups uniquely conserved across species. **(D-E) Clade E:** Specific receptor subregions of clade E. Highlighted receptors have some functional evidence (grey) or are de-orphanized with strong evidence (red). **(F) Clade J/K:** entire clade depicted, highlighting specific de-orphanized receptor groups that detect estrus cues (sulfated estrogens) in female urine.

#### Clade C

Clade C is the largest V1R clade in all non-commensal species examined in this study, exhibiting variable repertoire sizes suggestive of lineage-specific evolution (**Figure 4**). This inference is supported by the large numbers of combination-ID ortholog groups, gene duplicates, non-orthologous receptors, and house mouse-specific gene gains (**Figure 3**). The phylogenetic structure of clade C comprises three sub-clades. Interestingly, the non-orthologous receptors are largely clustered in one sub-clade, suggesting different rates of receptor evolution exist within clade C (**Figure 5B**). Two clade C receptors, *Vmn1r9* and *Vmn1r10*, have been implicated in pup odor detection in house mice (Isogai et al., 2018). However, these receptors also respond to female odors, and thus may detect chemosensory components of the nest environment (Isogai et al., 2018). These two receptors are part of a single receptor orthogroup (*Vmn1r9/10*) that is both orthologous and highly duplicated (**Figure 5B**). The sister group *Vmn1r7/8* exhibits a similar pattern of high orthology and abundant duplication (**Figure 5B**). Given the potential role of *Vmn1r9/10* receptors in pup odor detection, and the lineage-specific evolutionary patterns observed in *Vmn1r7/8* and *Vmn1r9/10*, these receptor groups are interesting candidates for future functional tests of their role in conspecific chemosignaling.

#### Clade D

Clade D exhibits a large skew in repertoire size within the house mouse (**Figure 4**), and has the most dramatic patterns of non-orthology of all V1R clades (**Figure 3B**). Nearly all reference-only V1Rs (50/53: 94%) are located in clade D, providing further support for a large recent gene expansion in the house mouse (**Figure 3B**). These receptors are similar in sequence and cluster together on chromosome 7, consistent with recent tandem gene duplication. While we did not find evidence for a comparably large expansion in the non-commensal species, we recover approximately twice as many clade D receptors in *M. spretus* relative to the other four species (**Figure 4**). It is possible that similar expansions exist in the other species that are not detected here. Additionally, clade D has a high proportion of non-orthologous receptors and gene duplicates. Given the evolutionary labile nature of clade D, there are a few rare conserved receptors that stand out: *V1rd19, Vmn1r179* and *Vmn1r172/173/174* (**Figure 5C**). While no functional data currently exists for these receptors, their distinct evolutionary pattern within clade D suggests they are under purifying selection, and are worthy of further investigation.

### Conserved Clades & Female-Specific Odor Detection

#### Clades E & F

A subset of V1R clades are highly conserved, and thus good targets for uncovering receptors with conserved olfactory functions. Clades E and F are characterized by high orthology (**Figure 3B**), long internal branch lengths and short terminal branch lengths, suggestive of old gene duplications maintained within the *Mus* lineage (**Additional File 3: Figure S3**). In contrast, very few recent gene duplications are detected (**Figure 3C, D**). A subset of 5 clade E receptors are important for the detection of female-specific urine odors in house mice (Haga-Yamanaka et al., 2014). Two clade E sub-regions containing all 5 receptor groups are shown in **Figure 5D, E**; those with the strongest support for female odor detection are highlighted in red (*Vmn1r69* and *Vmn1r185;* Haga-Yamanaka et al., 2014*). Vmn1r68* and *Vmn1r69* are sister to each other in the gene tree and are highly orthologous, however, *Vmn1r69* has no orthologs detected among the more basal species (*M. caroli* and *M. pahari*; **Figure 5D**). It is plausible that *Vmn1r69* is the result of a gene duplication event preceding the divergence of the four more derived species (**Figure 5D**), providing enhanced specificity or sensitivity toward female-specific urine odors. The second clade E sub-region contains receptors: *Vmn1r184, Vmn1r185*, and *Vmn1r71. Vmn1r184* and *Vmn1r185* are sister receptor groups, in which *Vmn1r185* is highly orthologous and *Vmn1r184* appears to be the result of a recent duplication event (**Figure 5E**). Interestingly, *Vmn1r184* is detected in only the house mouse and *M. spicilegus* (**Figure 5E**). Furthermore, *M. spicilegus* has evidence for a species-specific *Vmn1r184* duplicate, and has an absence of *Vmn1r185* expression (**Figure 5E**). The distinct expression pattern of *Vmn1r184* in *M. spicilegus* is noteworthy given this species’ unique social structure, which includes cooperative behaviors and social monogamy (Tong & Hoekstra, 2012). In comparison, *Vmn1r71* is highly orthologous (**Figure 5E**), but displays remarkable transcriptional variability, most of which is located at either the C-terminus or N-terminus regions of the protein (**Additional File 3: Figure S3**). Broadly, clades E and F display clade-wide patterns of conservation, with interesting lineage-specific receptor evolution within clade E surrounding sex-specific chemosignaling.

### Clade J/K Evolution & Detecting Estrus Cues

Clade J/K is both the most orthologous clade and boasts one of the highest proportions of gene duplicates (**Figure 3**). Thus this clade encompasses a unique mixture of conservation and divergence, in which there is very little gene loss but gene gains are abundant (**Figures 3 & 5F**). Clade J/K is also the only clade for which a large number of the receptors have known ligands. In the house mouse, two of the four J/K receptors (*Vmn1r85* and *Vmn1r89*) have been shown to detect estrus cues (i.e. sulfated estrogens) in female urine (Haga-Yamanaka et al., 2014; Isogai et al., 2011). The *Vmn1r89* receptor group has evidence for short and long transcript types across *Mus* species (**Additional File 3: Figure S4**). Many species have only one form detected, however, the house mouse and *M. spretus* express both forms as transcript variants of the same gene. While *M. pahari* appears to have distinct genes generating these two forms (**Additional File 3: Figure S4**). The widespread detection of both transcript types, suggests they may facilitate the detection of distinct ligand (i.e. sulfated estrogen) features. This is particularly compelling given that in the house mouse, *Vmn1r89*-expressing VSNs detect multiple sulfated estrogen molecules and are more broadly tuned than *Vmn1r85*-expressing VSNs (Haga-Yamanaka et al., 2014). In comparison, the *Vmn1r85* receptor group is highly conserved among the 3 *Mus* species most closely related to the house mouse (**Figure 5F**), with the majority of substitutions concentrated in *M. caroli* and *M. pahari* (**Additional File 3: Figure S5**). For both *Vmn1r85* and *Vmn1r89*, the highest proportion of amino acid site changes detected across species occurs in extracellular regions (**Additional File 3: Figure S6)**. The trend towards a higher rate of extracellular substitutions is consistent with a prior analysis of molecular evolution in 22 V1Rs, which demonstrated that most sites with evidence for positive selection are located in extracellular motifs (Emes et al., 2004).

## DISCUSSION

### V1R clades are characterized by distinct evolutionary trajectories

The complexity of the chemical environment presents unique evolutionary challenges. In addition to detecting a vast range of chemical stimuli, olfactory systems must flexibly adapt to novel environments and social contexts. One of the primary mechanisms of chemoreceptor evolution is through gene birth-and-death, mediated by duplication events and pseudogenization (Grus et al., 2005; Grus & Zhang, 2004; Silva & Antunes, 2017). Across divergent mammalian species, VRs have been shown to be fast-evolving with high gene turnover and lineage-specific clades, compared to the more conserved and largely orthologous ORs (Grus & Zhang, 2008). This has led to the hypothesis that ORs are broadly-tuned generalists, and VRs are more narrowly-tuned specialists (Grus & Zhang, 2008). Furthermore, olfactory specialization is hypothesized to occur through selection on distinct receptor subfamilies (Silva & Antunes, 2017). In this manner, receptor families may expand or contract in a lineage-specific fashion, and receptors in each family may become more diverse or conserved. Here, we identify distinct patterns of evolution among *Mus* V1R clades, consistent with a model of subfamily-specific selection. Some V1R clades have evidence for high gene turnover, while others are conserved across species. Prior research has generated controversy over what evolutionary forces mediate V1R evolution. Some studies detect evidence of positive selection and lineage-specific pseudogenization, while another study finds evidence for genetic drift and negative selection (Emes et al., 2004; Kurzweil et al., 2009; Park et al., 2011). Our data suggest that these seemingly contradictory results are not mutually exclusive. Depending on the subfamily of receptors examined, one could detect very different evolutionary patterns. This creates a functional framework in which to examine subsets of V1Rs, as receptor evolution is sculpted by the identity of their ligands.

### V1R gene gains and losses

Our results support the gene birth-and-death model of V1R evolution, exemplified by the variable patterns of orthology, gene duplicates, and sequence diversity observed across clades. However, while gene gains appear abundant across *Mus* species, gene losses are infrequent. A reliable estimate of V1R gene loss is restricted to the house mouse, due to constraints of V1R recovery among the other *Mus* species sequenced. Nevertheless, across all V1R clades only 10 gene losses are detected in the house mouse. This stands in contrast to a previous study examining the microevolution of V1Rs among *Mus musculus* subspecies, which detected a high frequency of null alleles (Park et al., 2011). We also identify a novel clade, which appears to have undergone pseudogenization in house mice and in rats. This is intriguing given the unique commensal status of house mice and rats relative to the other *Mus* species examined. As such, the clade N loss may reflect an adaptation to their unique commensal ecology. On the other hand, functional duplicates are plentiful. The most striking example is in the house mouse, in which clade D appears to have undergone a large species-specific gene expansion. This suggests that in the *Mus* genus, or perhaps solely within *Mus musculus* subspecies, expansion of the V1R family is ongoing.

### Patterns of receptor expression and evolution

Prior studies have examined expression levels of V1Rs across sexes and strains within the house mouse (Duyck et al., 2017; Ibarra-Soria et al., 2014). Interestingly, the expression patterns of V1R clades in the house mouse are often tightly correlated with evolutionary patterns observed across species. V1R clades that are highly conserved, are also the most highly expressed in the house mouse (clades: E, F and J/K; Duyck et al., 2017; Ibarra-Soria et al., 2014). In contrast, clades displaying rapid and species-specific evolution overlap with clades that include differentially expressed receptors among different house mouse strains (clades C and D; Duyck et al., 2017). Together, these two lines of evidence suggest that clades C and D are involved in species-specific chemical detection, while clades E, F and J/K likely perform crucial and conserved functions across mouse species. The obvious confound, however, is that clades with higher levels of expression may appear more orthologous due to high transcript recovery.

### Patterns of receptor evolution and function

Only a handful of V1Rs have known ligands. However, it has become increasingly clear that a critical function of the VNO involves detecting heterospecific odors, such as predator cues (Ben-Shaul et al., 2010; Isogai et al., 2011; Papes et al., 2010). V1Rs tuned to detecting broad classes of predator cues (e.g. birds of prey, snakes or mammals) may be conserved across mouse species. In particular, clade F has been implicated in detecting mammalian predator cues (Isogai et al., 2011). The broad-scale patterns of conservation observed in clade F are consistent with the maintenance of a similar key function, such as the detection of predator odor cues with shared ligands.

Chemical signaling is critical to social and reproductive interactions across a wide variety of mammalian species, including mice. One of the best described olfactory communication systems exists in house mouse urine scent marks (Hurst et al., 2001; Kaur et al., 2014; Sheehan et al., 2016). House mice secrete proteins in their urine (major urinary proteins, MUPs) that facilitate pheromonal communication and individual recognition (Hurst et al., 2001; Kaur et al., 2014; Roberts et al., 2012; Sheehan et al., 2016). MUPs act as transport vessels for the slow-release of volatile compounds detected by V1Rs (Bacchini, et al., 1992; Boschat et al., 2002; Novotny et al., 1999; Roberts et al., 2018). As these protein ligands vary considerably across *Mus* species, their corresponding volatiles likely shift as well (Roberts et al., 2018; Sheehan et al., 2019). As a result, clades such as C, D and H, exhibiting highly species-specific evolution may be good targets for the detection of social cues.

Mounting evidence suggests that V1Rs are crucial for detecting sex-specific cues as well as the physiological status of conspecifics (Celsi et al., 2012; Haga-Yamanaka et al., 2014; Isogai et al., 2011; Nodari et al., 2008). A subset of clade E receptors respond to female-specific urine ligands, thus clade E conservation may be tied to detecting conspecific sex cues (Haga-Yamanaka et al., 2014). Clade D has also been implicated in detecting female odors (Isogai et al., 2011). However, the activation of clade D is quite specific to *Vmn1r167* (Isogai et al., 2011). Interestingly, *Vmn1r167* contains one of the largest species-specific (*M. spicilegus*) gene duplications and is only detected in *M. spicilegus* and the house mouse. Therefore, *Vmn1r167* may play an important derived role in female odor detection.

Previous work demonstrated that V1Rs are strongly activated by sulfated steroids, and up to 80% of ligands detected in female urine may be sulfated steroids (Nodari et al., 2008). Clade J/K has been shown to play an important role in detecting sulfated estrogen molecules (Haga-Yamanaka et al., 2014). As such, the pattern of conservation in clade J/K may reflect a crucial role for these receptors in discerning information about the internal state of conspecifics, particularly female reproductive state.

## CONCLUSIONS

Understanding the evolutionary dynamics of the vomeronasal system reveals important properties of chemosensory evolution, as well as the functional roles of different receptors. In generating near-complete V1R repertoires for 5 *Mus* species, we find evidence for previously described patterns of high gene turnover observed among divergent species. However, by examining the evolutionary relationships of V1Rs across the *Mus* genus, we find that distinct receptor lineages have experienced different evolutionary trajectories. Thus, clade-level evolution is critical to understanding the chemosensory adaptions of species to their diverse chemical environments. Furthermore, the evolutionary patterns of V1Rs observed supports the proposition that the detection of physiological status and female-specific cues may be an important role of V1R chemosensation (Celsi et al., 2012; Haga-Yamanaka et al., 2014; Isogai et al., 2011; Nodari et al., 2008). Ultimately, these results provide a key foundation for future functional studies of V1Rs.

## METHODS

### Animal strains and tissues

Mouse strains for *M. caroli* (CAR: RBRC00823) and *M. spicilegus* (ZBN/Ms: RBRC00661) were obtained from RIKEN BioResource Center (Japan). *M. pahari* (PAH/EiJ) was obtained from The Jackson Laboratory (Bar Harbor, ME). All strains were maintained in an Animal Care facility at Cornell University with a 14:10 shifted light:dark cycle, and provided food and water *ad libitum*. Experimental protocols were approved by the Institutional Animal Care and Use Committee (IACUC: Protocol #2015-0060), and were in compliance with the NIH Guide for Care and Use of Animals. VNOs (stored in RNALater) for *M. macedonicus* (XBS) and *M. spretus* (SFM) were obtained from the Campbell Lab at Oklahoma State University (OSU). Mice at OSU were maintained on a 12:12 light:dark cycle and provided with food and water *ad libitum*. Live animal work at OSU was approved by the IACUC under protocol # AS-1-41.

### Illumina RNA library preparation & sequencing

VNO epithelia were dissected from at least one male and one female from each species and subsequently pooled to obtain V1R repertoires unbiased to a particular sex, except for the HiSeq dataset. Total RNA was extracted from VNO tissues using the Qiagen RNeasy kit, and subsequently quantified using QuBit Fluorometric Quantitation. RNA sequencing libraries were generated using the NEBNext Ultra RNA Library Prep Kit for Illumina (NEB #E7530). NEBNext Poly(A) mRNA Magnetic Isolation Module (NEB #E7490) was used for RNA Isolation. Sequences were indexed using the NEBNext Multiplex Oligos for Illumina (Dual Index Primers Set 1, NEB #E7600). A series of sequencing runs were performed on Illumina and PacBio platforms. The VNO libraries for strains ZRU, XBS and CAR were sequenced as 300 bp paired-end reads on Illumina MiSeq platform through the Biotechnology Resource Center (Institute of Biotechnology) at Cornell University. Additional VNO RNA libraries for strains ZRU, XBS, CAR and PAH were sequenced as 150 bp paired-end reads on Illumina NextSeq 500 platform through the Biotechnology Resource Center (Institute of Biotechnology) at Cornell University. A series of 15 SFM female VNO samples were sequenced as 125 bp paired-end reads on Illumina HiSeq 2500 platform at Novogene (Sacramento, CA). An additional round of long-read sequencing PacBio Isoseq libraries were generated and sequenced for SFM, using male and female VNOs to ensure the species-wide V1R repertoire was captured.

### Transcript processing and assembly

FastQC reports were generated for each sample to ensure sequencing quality (Andrews, 2010). Trimmomatic was used to clean the raw reads (Bolger et al., 2014). The trimmed read files were concatenated for each species across the different Illumina sequencing runs. rnaSPAdes was used to generate *de novo* transcriptome assemblies for each species’ concatenated RNA sequencing dataset (Bushmanova et al., 2018). Transrate and rnaQUAST were used for assembly quality assessment (Bushmanova et al., 2016; Smith-Unna et al., 2016). Other assemblers were tested (e.g. Trinity), however rnaSPAdes consistently assembled longer reads and more VRs were recovered from these assemblies.

### Isoseq library preparation and consensus assemblies

Isoseq was used to generate long-read sequences for the VNO from *Mus spretus* at the Arizona Genome Institute. We sequenced 4 different library sizes 0.8-1.6kb (x3 smartcells), 1.3-2.6kb (x2 smartcells), 2.2-3.7kb (x2 smartcells) and >3.0 kb (x2 smartcells) generating a total of 19GB of raw data. These data were run through the Pacbio smrtpipe version 2.3 by the Arizona Genome Institute, generating polished high consensus sequences, which we analyzed further for V1R genes.

### Identification of V1R sequences

The Ensembl reference annotation (version 94) of the mouse reference genome (GRCm38.p6) was used to download all known sequences for *V1rs*. These reference sequences were used in a series homology-based searches (blastn, blastx and tblastn) to identify putative *V1rs* in the RNA transcript assemblies for each mouse species. GetORF (http://emboss.sourceforge.net/apps/cvs/emboss/apps/getorf.html) was then used to identify open reading frames (ORFs) among the putative *V1r* dataset, using a well-defined *V1r* gene model (Ibarra-Soria et al., 2014). Dedupe was used to remove exact duplicate DNA sequences and containment DNA sequences within this refined ORF dataset. DNA sequences were translated into corresponding peptide sequences using GetORF. MAFFT v. 7 (https://mafft.cbrc.jp/alignment/software/) was used to align the peptide *V1r* sequences for each species, and sequences with less than 30% identity with the entire *V1r* group for a given species were eliminated (Katoh & Standley, 2013). While this pipeline was designed to identify functional *V1r* genes, given the abundance of *V1r* pseudogenes and the incomplete genome annotations for many of these species, some pseudogenes may inadvertently be included in these analyses.

### V1R annotation and identification of orthologous receptors

Putative *V1rs* were first annotated based on homology to the mouse reference genome. If multiple transcripts were *most similar* to a specific reference V1R gene (e.g., *Vmn1r30*), these transcripts were annotated with this same gene ID, and distinguished with unique numbers following the gene ID (e.g., *Vmn1r30.1* and *Vmn1r30.2*). Some annotations based on homology and their orientation within the gene tree did not always perfectly match due to the effects of gene duplications and losses at varying points in the *Mus* phylogeny. As such, some V1R annotations were adjusted upon analysis of the phylogenetic relationships of receptor sequences within the maximum-likelihood gene tree. The most important criteria for determining orthologous receptor groups was the relative orientation of all 6 species, under the general rule that the receptor phylogeny should recapitulate the species phylogeny. Using the annotation system of the reference genome meant that some gene duplications with distinct reference annotations were included in the same receptor ortholog group. Thus, some orthologous receptor groups were annotated with combination-IDs (e.g. *Vmn1r25/30*). Furthermore, a proportion of receptors from each of the five sequenced non-reference species were non-orthologous in that they did not fall into any particular ortholog group, but were basal to multiple groups or to several reference genes. These non-orthologous receptors were annotated based on the genes they were basal to, either as a combination-ID (e.g. *Vmn1r90/168/177*) or in the format “basalgeneID” if the number of gene IDs exceeded three (the lowest gene ID number was used). Thus V1R orthologs were identified using both sequence homology and phylogenetic relationships among receptors for all 6 species. Additionally, V1Rs that have evidence for gene losses in the house mouse (and corresponding pseudogenes) could not be annotated with the pseudogene ID, as often there are functional V1Rs with the same ID number. As a result, all gene losses in the house mouse detected in multiple non-commensal species are provided a new gene ID that does not overlap with any existing gene numbers. The annotated *V1r* coding sequences for all 5 non-commensal *Mus* species sequenced are provided in **Additional Files 4-8**.

### Phylogenetic analysis

All *V1r* peptide sequences for all 6 *Mus* species were aligned in MAFFT v. 7 (**Additional File 9**) (Katoh & Standley, 2013). Phylogenetic relationships were inferred using RAxML v. 8, which was used to generate a maximum likelihood gene trees (based on peptide sequences) with 1000 replicates of bootstrapping (**Additional File 1**) (Stamatakis, 2014). Trees were visualized in FigTree v1.4.3 (http://tree.bio.ed.ac.uk/software/) (Rambaut, 2016). A few traditionally separated clades were combined due to a lack of clear clade separation when viewed across all 6 *Mus* species (clades: A/B and J/K) (Rodriguez et al., 2002).

### Estimating gene duplicates

The well-characterized V1R repertoire of the reference genome was used to make estimates about which sequenced V1R transcripts are putative transcript variants or gene duplications within a given ortholog group (**Additional File 2**). Out of all the V1R transcripts in the reference, 55% code for the same peptide sequence, while 9.4% encode different peptides. Among the transcript variants encoding different peptides, sequence variation consists of either shorter sequences (i.e. only one exon is present) or variation at the ends of the transcript surrounding regions with gaps in pairwise alignments. We classified any V1R transcripts that (1) code for the same peptide, (2) whose variation consists of shortened coding sequences (i.e. only one exon is present), (3) whose variation falls at the ends of the transcript, or (4) whose variation falls near gaps in pairwise alignments, as putative transcript variants. Transcripts classified as putative gene duplicates were only those transcripts with at least one amino acid change central to the transcript, and not surrounded by gaps in pairwise alignments. This was only observed among different genes in the reference, never among transcript variants. Pairwise alignments were performed using EMBOSS Needle (https://www.ebi.ac.uk/Tools/psa/emboss_needle/). Due to the dynamic nature of V1R evolution and the incomplete V1R repertoires recovered for each species, duplications aren’t examined based on whether they are shared among species or are species-specific. Rather, duplications within ortholog groups are treated independently for each species.

### Motif prediction and mutation analysis

MAFFT v. 7 was used to align the orthologous receptor sequences (Katoh & Standley, 2013). TMHMM v. 2.0 (http://www.cbs.dtu.dk/services/TMHMM/) was used to predict the locations of transmembrane helices, and which V1R protein regions are intracellular versus extracellular (Krogh et al., 2001). For both *Vmn1r85* and *Vmn1r89* receptor groups, transcripts were aligned, transmembrane regions predicted, and sites with amino acid differences were identified across species (**Figures S4 & S5**). These amino acid differences are shown in protein schematics for each receptor (**Figure S6**). Four types of amino acid sites were characterized: (1) sites with an amino acid difference present in a single species, (2) sites with distinct amino acid differences in two different species, (3) sites with an amino acid difference shared between two to three species, (4) highly variable sites, in which amino acid differences suggest dynamism across the phylogenetic history of the genus (**Figure S6**). To examine *Vmn1r89* amino acid differences across species, the short transcriptional variants were excluded (**Figure S4**).

## Supporting information

Additional File 1. V1R gene tree

Additional File 2. Mus species V1R duplicates and trascript variants

Additional File 4. M. spicilegus V1R sequences

Additional File 5. M. macedonicus V1R sequences

Additional File 6. M. spretus V1R sequences

Additional File 7. M. caroli V1R sequences

Additional File 8. M. pahari V1R sequences

Additional File 9. V1R peptide sequence alignment

## ADDITIONAL FILES

**Additional File 1:** Maximum likelihood gene tree of all 6 *Mus* species’ V1R peptide sequences. (txt) **Additional File 2:** Categorization of V1R transcripts as either putative transcript variants or gene duplicates for the 5 non-commensal *Mus* species sequenced. (xlsx)

**Additional File 3: Table S1.** V1Rs with evidence for conservation (orthology and sequence identity) or gene expansions (across species or species-specific). **Figure S1.** Number of V1R genes with splice variants in *M. m. domesticus* and *M. spretus*. **Figure S2**. Novel clade “N”: *Vmn1r248* and *Vmn1r249.* **Figure S3. Left:** V1R gene tree clades E and F, displaying long internal branch lengths and short terminal branch lengths. ***Right***: Multiple alignments of *Vmn1r69* and *Vmn1r71* peptide sequences. **Figure S4.** Alignment and pairwise comparisons of *Vmn1r89* peptide sequences. **Figure S5.** Alignment and pairwise comparisons of *Vmn1r85* peptide sequences. **Figure S6.** Amino acid site changes in clade J/K receptors: *Vmn1r89* and *Vmn1r85*. (docx)

**Additional File 4:** *M. spicilegus* coding sequences of *V1r* genes expressed in the VNO. Gene annotations are abbreviated and contain species identifier “z”: *Vmn1r137* as 137-z. (fasta)

**Additional File 5:** *M. macedonicus* coding sequences of *V1r* genes expressed in the VNO. Gene annotations are abbreviated and contain species identifier “x”: *Vmn1r137* as 137-x. (fasta)

**Additional File 6:** *M. spretus* coding sequences of *V1r* genes expressed in the VNO. Gene annotations are abbreviated and contain species identifier “s”: *Vmn1r137* as 137-s. (fasta)

**Additional File 7:** *M. caroli* coding sequences of *V1r* genes expressed in the VNO. Gene annotations are abbreviated and contain species identifier “c”: *Vmn1r137* as 137-c. (fasta)

**Additional File 8:** *M. pahari* coding sequences of *V1r* genes expressed in the VNO. Gene annotations are abbreviated and contain species identifier “p”: *Vmn1r137* as 137-p. (fasta)

**Additional File 9:** Multiple sequence alignment of all 6 *Mus* species’ V1R peptide sequences. (fasta)

## LIST OF ABBREVIATIONS

OR: main olfactory receptor
VR: vomeronasal receptor
V1R: vomeronasal type 1 receptor
V2R: vomeronasal type 2 receptor
VNO: vomeronasal organ
Orthogroup: orthologous receptor group.

## ACKNOWLEDGEMENTS

The *M. macedonicus* (XBS), *M. spicilegus* (ZRU), and *M. spretus* (SFM) strains were originally developed by the Wild Mouse Genetic Repository (University of Montpellier). We thank Sara Miller for assistance with bioinformatic analyses.

## AUTHORS’ CONTRIBUTIONS

C.H.M. generated samples and sequence data, analyzed the data, and wrote the initial manuscript. M.J.S. and C.H.M. conceived and designed the experiment. P.C. contributed samples and sequencing data. All authors contributed to the editing of the manuscript.

## AVAILABILITY OF DATA AND MATERIALS

Newly sequenced vomeronasal organ transcriptomes supporting the conclusions of this article are available in the NCBI Short Read Archive under BioProject PRJNA596328. Sequences and datasets used for analyses are provided in additional files included within the article.

## CONSENT OF PUBLICATION

Not applicable.

## REFERENCES

Bacchini, A., Gaetani, E., & Cavaggioni, A. (1992). Pheromone binding proteins of the mouse, Mus musculus. Experientia, 48, 419–421.

Ben-Shaul, Y., Katz, L. C., Mooney, R., & Dulac, C. (2010). In vivo vomeronasal stimulation reveals sensory encoding of conspecific and allospecific cues by the mouse accessory olfactory bulb. Proceedings of the National Academy of Sciences, 107(11), 5172–5177.

Bolger, A. M., Lohse, M., & Usadel, B. (2014). Trimmomatic: A flexible trimmer for Illumina sequence data. Bioinformatics, 30(15), 2114–2120.

Boschat, C., Pélofi, C., Randin, O., Roppolo, D., Lüscher, C., Broillet, M. C., & Rodriguez, I. (2002). Pheromone detection mediated by a V1r vomeronasal receptor. Nature Neuroscience, 5(12), 1261–1262.

Brignall, A. C., & Cloutier, J. F. (2015). Neural map formation and sensory coding in the vomeronasal system. Cellular and Molecular Life Sciences, 72(24), 4697–4709.

Bushmanova, E., Antipov, D., Lapidus, A., & Przhibelskiy, A. D. (2018). rnaSPAdes: a de novo transcriptome assembler and its application to RNA-Seq data. BioRxiv, 420208.

Bushmanova, E., Antipov, D., Lapidus, A., Suvorov, V., & Prjibelski, A. D. (2016). RnaQUAST: A quality assessment tool for de novo transcriptome assemblies. Bioinformatics, 32(14), 2210–2212.

Celsi, F., D’Errico, A., & Menini, A. (2012). Responses to Sulfated Steroids of Female Mouse Vomeronasal Sensory Neurons. Chemical Senses, (37), 849–858.

Chamero, P., Marton, T. F., Logan, D. W., Flanagan, K., Cruz, J. R., Saghatelian, A., Cravatt, B.F., Stowers, L. (2007). Identification of protein pheromones that promote aggressive behaviour. Nature, 450(7171), 899–902.

Del Punta, K., Leinders-Zufall, T., Rodriguez, I., Jukam, D., Wysocki, C. J., Ogawa, S., Zufall, F., Mombaerts, P. (2002). Deficient pheromone responses in mice lacking a cluster of vomeronasal receptor genes. Nature, 419(6902), 70–74.

Dulac, C., & Wagner, S. (2006). Genetic Analysis of Brain Circuits Underlying Pheromone Signaling. Annual Review of Genetics, 40(1), 449–467.

Dulac, C., & Axel, R. (1995). A novel family of genes encoding putative pheromone receptors in mammals. Cell, 83(2), 195–206.

Dulac, C., & Torello, A. T. (2003). Molecular detection of pheromone signals in mammals: from genes to behavior. Nature Reviews Neuroscience, 4, 551–562.

Duyck, K., DuTell, V., Ma, L., Paulson, A., & Yu, C. R. (2017). Pronounced strain-specific chemosensory receptor gene expression in the mouse vomeronasal organ. BMC Genomics, 18(1), 1–16.

Emes, R. D., Beatson, S. A., Ponting, C. P., & Goodstadt, L. (2004). Evolution and comparative genomics of odorant-and pheromone-associated genes in rodents. Genome Research, 14(4), 591–602.

Ferrero, D. M., Moeller, L. M., Osakada, T., Horio, N., Li, Q., Roy, D. S., Cicht, A., Spehr, M., Touhara, K., Liberles, S. D. (2013). A juvenile mouse pheromone inhibits sexual behaviour through the vomeronasal system. Nature, 502(7471), 368–371.

Grus, Wendy E., Shi, P., Zhang, Y., & Zhang, J. (2005). Dramatic variation of the vomeronasal pheromone receptor gene repertoire among five orders of placental and marsupial mammals. Proceedings of the National Academy of Sciences, 102(16), 5767–5772.

Grus, Wendy E., & Zhang, J. (2004). Rapid turnover and species-specificity of vomeronasal pheromone receptor genes in mice and rats. Gene, 340(2), 303–312.

Grus, Wendy E, & Zhang, J. (2008). Distinct Evolutionary Patterns between Chemoreceptors of 2 Vertebrate Olfactory Systems and the Differential Tuning Hypothesis. Molecular Biology and Evolution, 25(8), 1593–1601.

Haga-Yamanaka, S., Ma, L., He, J., Qiu, Q., Lavis, L. D., Looger, L. L., & Yu, C. R. (2014). Integrated action of pheromone signals in promoting courtship behavior in male mice. ELife, 2014(3), 1–19.

Haga, S., Hattori, T., Sato, T., Sato, K., Matsuda, S., Kobayakawa, R., Sakano, H., Yoshihara, Y., Kikusui, T., Touhara, K. (2010). The male mouse pheromone ESP1 enhances female sexual receptive behaviour through a specific vomeronasal receptor. Nature, 466(7302), 118–122.

He, J., Ma, L., Kim, S., Nakai, J., & Yu, C. R. (2008). Encoding gender and individual Information in the Mouse. Science, 320(5875), 535–538.

Hurst, J. L., Payne, C. E., Nevison, C. M., Marie, a D., Humphries, R. E., Robertson, D. H., Cavaggioni, A., Beynon, R. J. (2001). Individual recognition in mice mediated by major urinary proteins. Nature, 414(6864), 631–634.

Ibarra-soria, X., Levitin, M. O., Saraiva, L. R., & Logan, D. W. (2014). The Olfactory Transcriptomes of Mice, 10(9).

Isogai, Y., Si, S., Pont-Lezica, L., Tan, T., Kapoor, V., Murthy, V. N., & Dulac, C. (2011). Molecular organization of vomeronasal chemoreception. Nature, 478(7368), 241–245.

Isogai, Y., Wu, Z., Love, M. I., Ahn, M. H. Y., Bambah-Mukku, D., Hua, V., Farrell, K., Dulac, C. (2018). Multisensory Logic of Infant-Directed Aggression by Males. Cell, 175(7), 1827-1841.e17.

Jiao, H., Hong, W., Nevo, E., Li, K., & Zhao, H. (2019). Convergent reduction of V1R genes in subterranean rodents. BMC Evolutionary Biology, 19(176), 1–9.

Katoh, K., & Standley, D. M. (2013). MAFFT multiple sequence alignment software version 7: Improvements in performance and usability. Molecular Biology and Evolution, 30(4), 772–780.

Kaur, A. W., Ackels, T., Kuo, T. H., Cichy, A., Dey, S., Hays, C., Kateri, M., Logan, D. W., Marton, T. F., Spehr, M., Stowers, L. (2014). Murine pheromone proteins constitute a context-dependent combinatorial code governing multiple social behaviors. Cell, 157(3), 676–688.

Kimoto, H., Haga, S., Sato, K., & Touhara, K. (2005). Sex-specific peptides from exocrine glands stimulate mouse vomeronasal sensory neurons. Nature, 437(7060), 898–901.

Krogh, A., Larsson, B., Von Heijne, G., & Sonnhammer, E. L. L. (2001). Predicting transmembrane protein topology with a hidden Markov model: Application to complete genomes. Journal of Molecular Biology, 305(3), 567–580.

Kurzweil, V. C., Getman, M., Green, E. D., & Lane, R. P. (2009). Dynamic evolution of V1R putative pheromone receptors between Mus musculus and Mus spretus. BMC Genomics, 10, 1–11.

Lane, R. P., Young, J., Newman, T., & Trask, B. J. (2004). Species Specificity in Rodent Pheromone Receptor Repertoires. Genome Research, 14, 603–608.

Leinders-Zufall, T. (2000). Ultrasensitive pheromone detection by mammalian vomeronasal neurons. Letters to Nature, 405, 251–260.

Leinders-Zufall, T., Brennan, P., & Widmayer, P. (2004). MHC Class I Peptides as Chemosensory Signals in the Vomeronasal Organ. Science, 306, 1033–1038.

Leypold, B. G., Yu, C. R., Leinders-Zufall, T., Kim, M. M., Zufall, F., & Axel, R. (2002). Altered sexual and social behaviors in trp2 mutant mice. Proceedings of the National Academy of Sciences, 99(9), 6376–6381.

Liberles, S. D. (2014). Mammalian Pheromones. Annual Review of Physiology, 76(1), 151–175.

Meeks, J. P., Arnson, H. A., & Holy, T. E. (2010). Representation and transformation of sensory information in the mouse accessory olfactory system. Nature Neuroscience, 13(6), 723–730.

Meisami, E., & Bhatnagar, K. P. (1998). Structure and Diversity in Mammalian Accessory Olfactory Bulb. Microscopy Research and Technique, 43, 476–499.

Mombaerts, P. (2004). Genes and ligands for odorant, vomeronasal and taste receptors. Nature Reviews Neuroscience, 5(4), 263–278.

Nara, K., Saraiva, L. R., Ye, X., & Buck, L. B. (2011). A Large-Scale Analysis of Odor Coding in the Olfactory Epithelium. Journal of Neuroscience, 31(25), 9179–9191.

Nodari, F., Hsu, F., Fu, X., Holekamp, T. F., Kao, L., Turk, J., & Holy, T. E. (2008). Sulfated Steroids as Natural Ligands of Mouse Pheromone-Sensing Neurons. Journal of Neuroscience, 28(25), 6407–6418.

Novotny, M., Jemiolo, B., Harvey, S., Wiesler, D., & Marchlewska-koj, A. (1986). Adrenal-Mediated Endogenous Metabolites Inhibit Puberty in Female Mice. Science, 231(4739), 722–725.

Novotny, M. V, Ma, W., Wiesler, D., & Zídek, L. (1999). Positive identification of the puberty-accelerating pheromone of the house mouse: the volatile ligands associating with the major urinary protein. Proceedings of the Royal Society of London, Series B, 266, 2017–2022.

Orikasa, C., Kondo, Y., Katsumata, H., Terada, M., Akimoto, T., Sakuma, Y., & Minami, S. (2017). Physiology & Behavior Vomeronasal signal de fi ciency enhances parental behavior in socially isolated male mice. Physiology & Behavior, 168, 98–102.

Papes, F., Logan, D. W., & Stowers, L. (2010). The Vomeronasal Organ Mediates Interspecies Defensive Behaviors through Detection of Protein Pheromone Homologs. Cell, 141(4), 692–703.

Paradis, E., & Schliep, K. (2019). ape 5.0: an environment for modern phylogenetics and evolutionary analyses in R. Bioinformatics (Oxford, England), 35(3), 526–528.

Park, S. H., Podlaha, O., Grus, W. E., & Zhang, J. (2011). The microevolution of V1r vomeronasal receptor genes in mice. Genome Biology and Evolution, 3(1), 401–412.

Pocock, M. J. O., Searle, J. B., & White, P. C. L. (2004). Adaptations of animals to commensal habitats:population dynamics of house mice Mus musculus domesticus on farms. Journal of Animal Ecology, 73, 878–888.

Powers, J. B., & Winans, S. S. (1975). Vomeronasal Organ: Critical Role in Mediating Sexual Behavior of the Male Hamster. Science, 961–964.

Restrepo, D., Arellano, J., Oliva, A. M., Schaefer, M. L., & Lin, W. (2004). Emerging views on the distinct but related roles of the main and accessory olfactory systems in responsiveness to chemosensory signals in mice. Hormones and Behavior, 46, 247–256.

Roberts, S. A., Davidson, A. J., McLean, L., Beynon, R. J., & Hurst, J. L. (2012). Pheromonal induction of spatial learning in mice. Science, 338(6113), 1462–1465.

Roberts, S. A., Prescott, M. C., Davidson, A. J., McLean, L., Beynon, R. J., & Hurst, J. L. (2018). Individual odour signatures that mice learn are shaped by involatile major urinary proteins (MUPs). BMC Biology, 16(1), 1–19.

Rodriguez, I. (2016). Vomeronasal Receptors: V1Rs, V2Rs, and FPRs. Chemosensory Transduction: The Detection of Odors, Tastes, and Other Chemostimuli. Elsevier Inc.

Rodriguez, I., Del Punta, K., Rothman, A., Ishii, T., & Mombaerts, P. (2002). Multiple new and isolated families within the mouse superfamily of V1r vomeronasal receptors. Nature Neuroscience, 5(2), 134–140.

Sheehan, M. J., Campbell, P., & Miller, C. H. (2019). Evolutionary patterns of major urinary protein scent signals in house mice and relatives. Molecular Ecology, 1–15.

Sheehan, M. J., Lee, V., Corbett-Detig, R., Bi, K., Beynon, R. J., Hurst, J. L., & Nachman, M. W. (2016). Selection on Coding and Regulatory Variation Maintains Individuality in Major Urinary Protein Scent Marks in Wild Mice. PLoS Genetics, 12(3), 1–33.

Shi, P., Bielawski, J. P., Yang, H., & Zhang, Y. P. (2005). Adaptive diversification of vomeronasal receptor 1 genes in rodents. Journal of Molecular Evolution, 60(5), 566–576.

Silva, L., & Antunes, A. (2017). Vomeronasal Receptors in Vertebrates and the Evolution of Pheromone Detection. Annual Review of Animal Biosciences, 5(1), 353–370.

Smith-unna, R., Boursnell, C., Patro, R., Hibberd, J. M., & Kelly, S. (2016). TransRate:reference-free quality assessment of de novo transcriptome assemblies. Genome Research, 26, 1134–1144.

Stamatakis, A. (2014). RAxML version 8: A tool for phylogenetic analysis and post-analysis of large phylogenies. Bioinformatics, 30(9), 1312–1313.

Stowers, L., Holy, T. E., Meister, M., Dulac, C., & Koentges, G. (2002). Loss of sex discrimination and male[ndash]male aggression in mice deficient for TRP2. Science, 295(February), 1493–1500.

Suzuki, H., Nunome, M., Kinoshita, G., Aplin, K. P., Vogel, P., Kryukov, A. P., Lin, M-L., Han, S-H.,Maryanto, I.,Tsuchiya, K., Ikeda, H., Yonekawa, H., Moriwaki, K. (2013). Evolutionary and dispersal history of Eurasian house mice Mus musculus clarified by more extensive geographic sampling of mitochondrial DNA. Heredity, 111(5), 375–390.

Tachikawa, K. S., Yoshihara, Y., & Kuroda, K. O. (2013). Behavioral transition from attack to parenting in male mice: A crucial role of the vomeronasal system. Journal of Neuroscience, 158(6), 5120–5126.

Tong, W., & Hoekstra, H. (2012). Mus spicilegus. Current Biology, 22(20), 858–859.

Wagner, S., Gresser, A. L., Torello, A. T., & Dulac, C. (2006). A Multireceptor Genetic Approach Uncovers an Ordered Integration of VNO Sensory Inputs in the Accessory Olfactory Bulb. Neuron, 50(5), 697–709.

Wynn, E. H., Sánchez-Andrade, G., Carss, K. J., & Logan, D. W. (2012). Genomic variation in the vomeronasal receptor gene repertoires of inbred mice. BMC Genomics, 13(1), 19–23.

Yang, H., Shi, P., Zhang, Y. P., & Zhang, J. (2005). Composition and evolution of the V2r vomeronasal receptor gene repertoire in mice and rats. Genomics, 86(3), 306–315.

Young, J. M., Massa, H. F., Hsu, L., & Trask, B. J. (2010). Extreme variability among mammalian V1R gene families. Genome Research, 20(1), 10–18.

