## Additional File 3. Supplemental Table S1 & Figures S1-S6 for "Distinct evolutionary trajectories of V1R clades across mouse species"

**Table S1. V1Rs with evidence for conservation (orthology and sequence identity) or gene expansions (across species or species-specific).** Species are indicated with letters: *M. m. domesticus* (m), *M. spicilegus* (z), *M. macedonics* (x), *M. spretus* (s), *M. caroli* (c), and *M. pahari* (p).

| Clade | Clade-Wide Evolutionary Pattern | Conserved: Highest Orthology | Conserved: % Seq ID | Multi-Species Expansions | Species-Specific Expansions | House Mouse Specific Expansions |
|---|---|---|---|---|---|---|
| **A/B** | Somewhat Dynamic | Vmn1r1, Vmn1r40, Vmn1r44, Vmn1r47/48 | Vmn1r51 | Vmn1r45, Vmn1r47/48 | Vmn1r44 (p), Vmn1r47/48 (s), Vmn1r50 (s) | Vmn1r52/V1ra8 |
| **C** | Dynamic | Vmn1r11, Vmn1r13, Vmn1r19, Vmn1r20/27, Vmn1r24, Vmn1r25/30, Vmn1r26, Vmn1r38/39 | --- | Vmn1r19, Vmn1r38/39, Vmn1r7/8, Vmn1r9/10 | Vmn1r19 (c), Vmn1r33 (s), Vmn1r36/37 (p), Vmn1r38/39 (z, s, p), Vmn1r7/8 (p) | --- |
| **D** | Dynamic | Vmn1r172/173/174, Vmn1r179, V1rd19 | --- | Vmn1r168/177, Vmn1r180 | Vmn1r167 (z), Vmn1r90 (s) | Vmn1r172/173/174, Vmn1r60/61, Vmn1r62/186, Vmn1r63/187, Vmn1r100/148, Vmn1r170/175, Vmn1r56, Vmn1r57, Vmn1r91, Vmn1r93-95, Vmn1r101, Vmn1r103, Vmn1r104, Vmn1r107, Vmn1r111-132, Vmn1r135, Vmn1r137-139, Vmn1r142, Vmn1r143, Vmn1r149, Vmn1r151, Vmn1r152, Vmn1r155, Vmn1r157-160, Vmn1r163, Vmn1r165, Vmn1r166, Vmn1r171, Vmn1r100/148 |
| **E** | Conserved | Vmn1r224, Vmn1r227, Vmn1r230, Vmn1r231, Vmn1r68, Vmn1r71 | Vmn1r224, Vmn1r227 | Vmn1r71 | --- | --- |
| **F** | Conserved | Vmn1r236, Vmn1r237 | --- | --- | --- | --- |
| **G** | Somewhat Conserved | Vmn1r73, Vmn1r74, Vmn1r79/2/3/238, Vmn1r84 | Vmn1r84 | --- | --- | Vmn1r2/3/79/238 |
| **H** | Dynamic | Vmn1r196, Vmn1r206/209, Vmn1r214 | Vmn1r197 | Vmn1r206/209, Vmn1r247 | Vmn1r203 (s), Vmn1r205 (s), Vmn1r206/209 (c, p) | --- |
| **I** | Somewhat Dynamic | Vmn1r192, Vmn1r193, Vmn1r194, Vmn1r218 | Vmn1r192, Vmn1r218 | Vmn1r247, Vmn1r192, Vmn1r193, Vmn1r202, Vmn1r216 | --- | --- |
| **J/K** | Mixed: Conserved & Dynamic | Vmn1r85, Vmn1r86, Vmn1r89 | --- | Vmn1r86/88 | Vmn1r86/88 (c) | --- |
| **L** | Conserved | --- | Vmn1r70 | --- | --- | --- |
| **N** | Mixed: Conserved & Dynamic | --- | --- | --- | --- | --- |

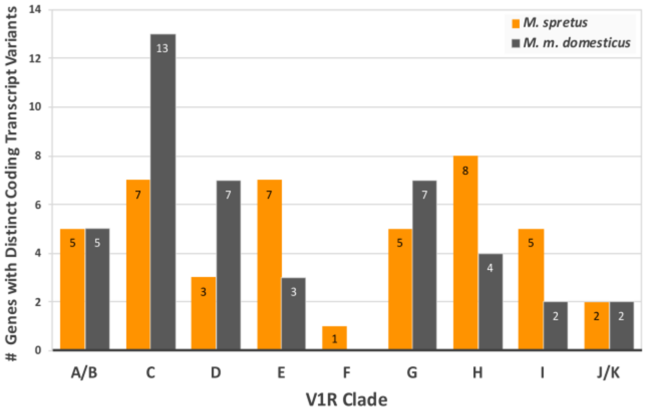


**Figure S1. Number of V1R genes with splice variants.** V1R genes by clade that express multiple coding transcripts with distinct peptide sequences in both the house mouse (grey) and *M. spretus* (orange).

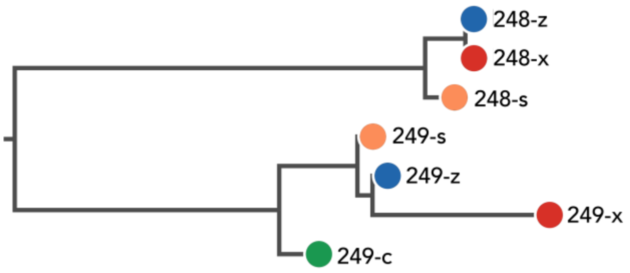


**Figure S2. Novel clade "N".** V1R gene tree clade N with genes *Vmn1r248* and *Vmn1r249*. *Mus* species are indicated with letter abbrevations and colors (*M. spicilegus*: "z" and blue; *M. macedonics*: "x" and red; *M. spretus*: "s" and orange; and *M. caroli*: "c" and green).

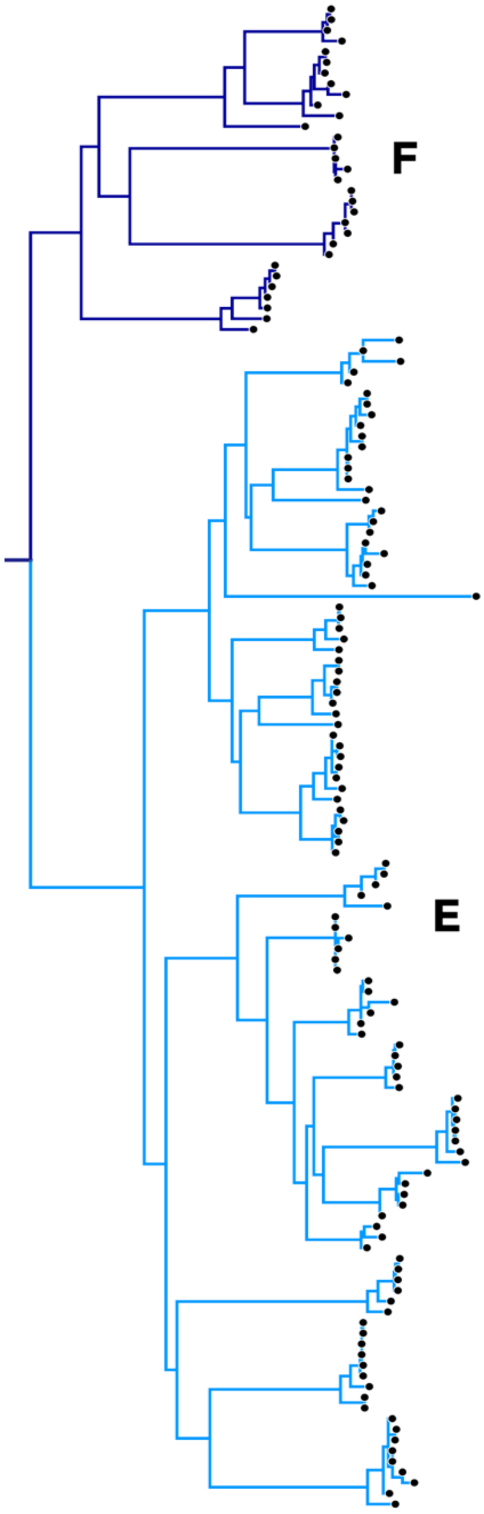


```
69.1-m      ------------------------------------------------------------
69.2-m      --------------------------------------------MGVATIKENQAQSKMD
69-s        MCTRTPCKYNSRCDRFSKNGHSFSSSYHLQGVLQCQCMGTGRQKMGVAIIKENQAQSKMD
69-z        ------------------------------------------------------------
69-x        ------------------------------------------------------------

69.1-m      ------------------------------------------------------------
69.2-m      FRNLAIKIIFLSQTTIGILGNFSLIIYYVVYYIVYTLKPTDLILMHLVTSNALMVLSTGV
69-s        FRNLAIKIIFLSQTTIGILGNFSLIIYYVVYSIDHTLKPTDLILMHLVTSNALMVLSTGV
69-z        ------------------------------------------------------------
69-x        --------------------------------------------MHLVTSNALMVLSTGV

69.1-m      ---MAAFGLTHLLNDLGCRLILYIQRVGRSISIGSTCLLSIFQAIIISHRESCYKGQKVK
69.2-m      PHTMAAFGLTHLLNDLGCRLILYIQRVGRSISIGSTCLLSIFQAIIISHRESCYKGQKVK
69-s        PHTMAAFGLTHLLNDLGCRLILYIQRVGRSISIGSTCLLSIFQAIIISHRESCYKGQKVK
69-z        ---MAAFGLTHLLNDLGCRLILYIQRVGRSISIGSTCLLSIFQAIIISHRESCYKGQKVK
69-x        PHTMAAFGLTHLLNDLGCRLILYIQRVGRSISIGSTCLLSIFQAIIISHWKSCYKGQKVK
               ********************************************** :*********

69.1-m      SAKYVGYSLSLLWVLHIFINFILFVDIFIKRNRNNMTRDRDFKYCYIVGQNEFSDLLYVA
69.2-m      SAKYVGYSLSLLWVLHIFINFILFVDIFIKRNRNNMTRDRDFKYCYIVGQNEFSDLLYVA
69-s        SAKYVGYSLSLLWVLHIFINFILFVDIFIKRNRNNMTRDRDFKYCYIVGQNEFSDLLYVA
69-z        SAKYVGYSLSLLWVLHIFINFILFVDIFIKRNRDNMTRDRDFKYCYIVGQNEFSDLLYVA
69-x        SAKYVGYSLSLLWVLHIFINFILFVDIFIKRNRDNMTRDRDFKYCYIVGQNEFSDLLYVA
            *********************************:**************************

69.1-m      LVVCPEILSSVLITYSSGSMIGILYRHKQRVQHIRSFHVSSRNSPEWRATQNILVLVSTF
69.2-m      LVVCPEILSSVLITYSSGSMIGILYRHKQRVQHIRSFHVSSRNSPEWRATQNILVLVSTF
69-s        LVVCPEILSSVLITYSSGSMIGILYRHKQRVQHIRSFHVSSRSSPEWRATQNILVLVSTF
69-z        LVVCPEILFSVLITYSSGSMIGILYRHKQRVQHIRSLHVSSRNSPEWRATQNILVLVSTF
69-x        LVVCPEILFSVLITYSSGSMIGILYRHKQRVQHIRSLHVSSRNSPEWRATQNILVLVSTF
            ******** ***************************:*****.*****************

69.1-m      LGFYTLSSILQGCIALQNNPNWWLLNINRLTSMCFPTFAPFVLMSHYSFMPKLSLTWIRN
69.2-m      LGFYTLSSILQGCIALQNNPNWWLLNINRLTSMCFPTFAPFVLMSHYSFMPKLSLTWIRN
69-s        LGFYTLSSILQGCIALLNNPNWWLLNINRLTSLCFPTFAPFVLMSHYSFMPKLSLTWIRN
69-z        LGFYTLSSILQGCIALLNNPNWWLLNINRLTSMCFPTFAPFVLMSHYSFMPKLSLTWIRN
69-x        LGFYTLSSILQGCIALLNNPNWWLLNINRLTSMCFPTFAPFVLMSHYSFMPKLSLTWIRN
            **************** ***************:***************************

69.1-m      KTS
69.2-m      KTS
69-s        KTS
69-z        KTS
69-x        KTS
            ***
```

```
71.1-m      -------------------------MDFWNLAIRIIFLLQTVIGILGNVFLIFYYLVLYY
71.2-m      ------------------------------------------------------------
71-x        -------------------------MDFWNLAIRIIFLSQTVTGTLGNVFLIFYYLVLYY
71.1-s      -------------------------MDFWNLAIRIIFLLQTVTGILGNVFLIFYYLVLYY
71.2-s      -------------------------MDFWNLAIRIIFLLQTVTGILGNVFLIFYYLVLYY
71.3-s      -------------------------MDFWNLAIRIIFLLQTVTGILGNVFLIFYYLVLYY
71.1-z      -------------------------MDFWNLEIRIIFLSQTVTGTLGNVFLIFYYLVLYY
71.2-z      -------------------------MDFWNLEIRIIFLSQTVTGTLGNVFLIFYYLVLYY
71-c        -------------------------MDFWNLAIRIIFLSQTATGILGNVFLIFYYLVLYY
71.1-p      -------------------------MDFWNLAIRIIFLSQTATGILGNFFLIFYYLTLYY
71.2-p      MLLTLFFWSYNFCGHPGKEIVLNDIMDLWNMAIRIIFLLQTTTGIFGNSSLILYYLICYY

71.1-m      RECTLKPTDLILMNLMAANLVILLSIGVPQKMAIWGLKEFLNHLECEILLFVQGFSRSMS
71.2-m      -------------------------------MAIWGLKEFLNHLECEILLFVQGFSRSMS
71-x        RECTLKPTDLILMNLMAANLVILLSIGVPQTMAIWGLKEFLNHLECEILLFVQGFSRSMS
71.1-s      RECTLKPTDLILMNLMAANLVILLSIGVPQTMAIWGLKEFLNHLECELLLFIQGFSRSMS
71.2-s      RECTLKPTDLILMNLMAANLVILLSIGVPQTMAIWGLKEFLNHLECELLLFIQGFSRSMS
71.3-s      RECTLKPTDLILMNLMAANLVILLSIGVPQTMAIWGLKEFLNHLECELLLFIQGFSRSMS
71.1-z      RECTLKPTDLILMNLMAANLVILLSIGVPQTMAIWGLKEFLNHLECEILLFIQGFSRSMS
71.2-z      RECTLKPTDLILMNLMAANLVILLSIGVPQTMAIWGLKEFLNHLECEILLFIQGFSRSMS
71-c        RECTLKPTDLILMNLLAANLVILLSIGVPQTMAIWGLKEFLNHLECELLLFIQGFSRSMS
71.1-p      RECTLKPTDLILMNLMAANLVILLAIGVPQTMAIWGLKEFLNHLECDLLLFIQGFGRSVS
71.2-p      KKYTLKPTDFILFHIMTANALIILSLGVPQTLSVWGLKEFLNNFGCDLLLYIQGFGRRVS
                                           :::********:: *::**::***.* :*

71.1-m      ICTICLLSVFQTLTISPKKFCWKYYKSNAAKYIGSCTSLLWVLHVLINFIFFLYTFTKRN
71.2-m      ICTICLLSVFQTLTISPKKFCWKYYKSNAAKYIGSCTSLLWVLHVLINFIFFLYTFTKRN
71-x        ICTICLLSVFQTLTISPKKFCWKYYKSKAAKYIGSCTSLLWVLHVLINFIFFLYTFIKRN
71.1-s      ICTICLLSVFQTLTISPKKFCWKYYKSKAAKYIGSCTSLLWVLHMLINFIFVLYTFIKRN
71.2-s      ICTICLLSVFQTLTISPKKFCWKYYKSKAAKYIGSCTSLLWVLHMLINFIFVLYTFIKRN
71.3-s      ICTICLLSVFQTLTISPKKFCWKYYKSKAAKYIGSCTSLLWVLHMLINFIFFLYTFIKRN
71.1-z      ICTICLLSVFQTLTISPKKFCWKYYKSNAAKYIGSCTSLLWVLHMLINFIFFLYTFIKRN
71.2-z      ICTICLLSVFQTLTISPKKFCWKYYKSNAAKYIGSCTSLLWVLHMLINFIFFLYTFIKRN
71-c        ICTICLLSVFQTLTISPKKFCWKYYKSKAAKYIGSCTSLLWVLHVLINFIFFLYTFIKRN
71.1-p      IYTICLLFVFQTVTISPKKFGWKYYKSKAVKHIGSCTSLLWVLPMLINFILFLYTFIKRN
71.2-p      ICMTCLLSIFQTITINPRKLCWKYHKVVASKVIGCSISLFWVLHMLINFIFFVYTSIKRN
            *   *** :***:**.*:*: ***:*  * * **.. **:*** :*****:.:**  ***

71.1-m      NTNVTRTRDFGYCSLIGSDDISDSLYAALVVCPEVFFSVLTALASVSMIVILYRHKKSVQ
71.2-m      NTNVTRTRDFGYCSLIGSDDISDSLYAALVVCPEVFFSVLTALASVSMIVILYRHKKSVQ
71-x        NTNVTRTRDFGYCSLIGSDDISDSLYAALVVCPEVFFSVLIAFTSVSMIVILYRHKKSVQ
71.1-s      NTNVTRTRDFGYCSLIGSDDISDSLYAALVVCPEVFFSVLTALASVSMIVILYRHKKRVQ
71.2-s      NTNVTRTRDFGYCSLIGSDDISDSLYAALVVCPEVFFSVLTALASVSMIVILYRHKKRVQ
71.3-s      NTNVTRTRDFGYCSLIGSDDISDSLYAALVVCPEVFFSVLTALASVSMIVILYRHKKRVQ
71.1-z      NRNVTRTRDFGYCSLIGSDDISDLLYAALVVCPEVFFSVLTAFTSVSMIVILYRHKKRVQ
71.2-z      NRNVTRTRDFGYCSLIGSDDISDLLYAALVVCPEVFFSVLTAFTSVSMIVILYRHKKRVQ
71-c        NTNVTRTRDFGYCSHTGSDDISDLIYAALVVCPEVFFSVLTVFTSVSMIVILYRHKKRVQ
71.1-p      NTNVTRIRDFGYCSLTGSDDISDLLYTVLVVCPEVFFSMLTALASVSMIVILYRHKKRVQ
71.2-p      RKNVTRIRDLGYCFLIGSDDISDFLYTALVVCPEVIFSVFIAWSSGSMIVILYRHKQRVQ
            . **** **:***   ******* :*:.*******:**:: . :* **********: **

71.1-m      HIRSTHGSSRYSPESRATQNILTVVSTFLAFCSLTTILRGFVALFHGHSWWLMNINHLTS
71.2-m      HIRSTHGSSRYSPESRATQNILTVVSTFLAFCSLTTILRGFVALFHGHSWWLMNINHLTS
71-x        HIRSTHGSSRYSPESRATQNILTVVSTFLAFCTLTSILRGCVALFHGHSWWLMNINHLTS
71.1-s      HIRSTHGSSRYSPESRATQNILTVVSTFLAFCTLTSILRGCVALFHGHSWWLMNINHLTS
71.2-s      HIRSTHGSSRYSPESRATQNILTVVSTFLAFCTL--------------------------
71.3-s      HIRSTHGSSRYSPESRATQNILTVVSTFLAFCTLTSILRGCVALFHGHSWWLMNINHLTS
71.1-z      HIRSTHGSSRYSPESRATQNILTVVSTFLAFCSLTTILRGFVALFHGHSWWLMNINHLTS
71.2-z      HIRSTHGSSRYSPESRATQNILTVVSTFLAFCSLTTILRGFVALFHGHSWWLMNINHLTS
71-c        HIRSTHGSSRYSPESRATQNILTVVSTFLAFCSLTTILRGCVALFHGHSWWLMNINHLTS
71.1-p      HIHSTHGSSQYSPESRATQSILTVVSTFLAFCTLTSILRGCVALFHGHSWWLMNINHLTS
71.2-p      YIHSSHRSCRCSTESRATQNILALLSTFLTCYTLSSILRGCIAFLHNHNWWLMNINHFIS
            :*:*:* *.: *.******.**:::****:   :*

71.1-m      LCFPTFAPFVLMSHYSIVSRLSLVWLRKKNIS------------------
71.2-m      LCFPTFAPFVLMSHYSIVSRLSLVWLRKKNIS------------------
71-x        LCFPTFAPFVLMSHYSIVSRLSLVWLRKK-ISHNSLIAQLFTHSTLIGQ
71.1-s      LCFPTFAPFVLMSHYSIVSRLSLVWLRKK-ISHNSLIAQLFTHS-----
71.2-s      -------------------------------------------------
71.3-s      LCFPTFAPFVLMSHYSIVSRLSLVWLKKKNIS------------------
71.1-z      LCFPTFAPFVLMSHYSIVSRLSLVWLRKK-ISHNSLIAQLFTHSSLIGQ
71.2-z      LCFPTFAPFVLMSHYSIVSRLSLVWLRKK-ISHNSLIAQLFTHS-----
71-c        LCFPTFAPFVLMSHYSIVSRLSLVWLRKKKIS------------------
71.1-p      LCFPTFSPFVLMSHYSIVSRLSLVCFRKKK-SHNSLIAQLFTHS-----
71.2-p      LCFPSFAPFVLISHYSMVSRFSLYWLRKKK-------------------
```

**Figure S3.** ***Left***: V1R gene tree clades E (light blue) and F (dark blue), displaying long internal branch lengths and short terminal branch lengths. ***Right***: Multiple alignment of *Vmn1r69* (top) and *Vmn1r71* (bottom) peptide sequences. *Mus* species indicated with letter abbrevations (*M. m. domesticus*: "m", *M. spicilegus*: "z", *M. macedonics*: "x", *M. spretus*: "s", *M. caroli*: "c" and *M. pahari*: "p").

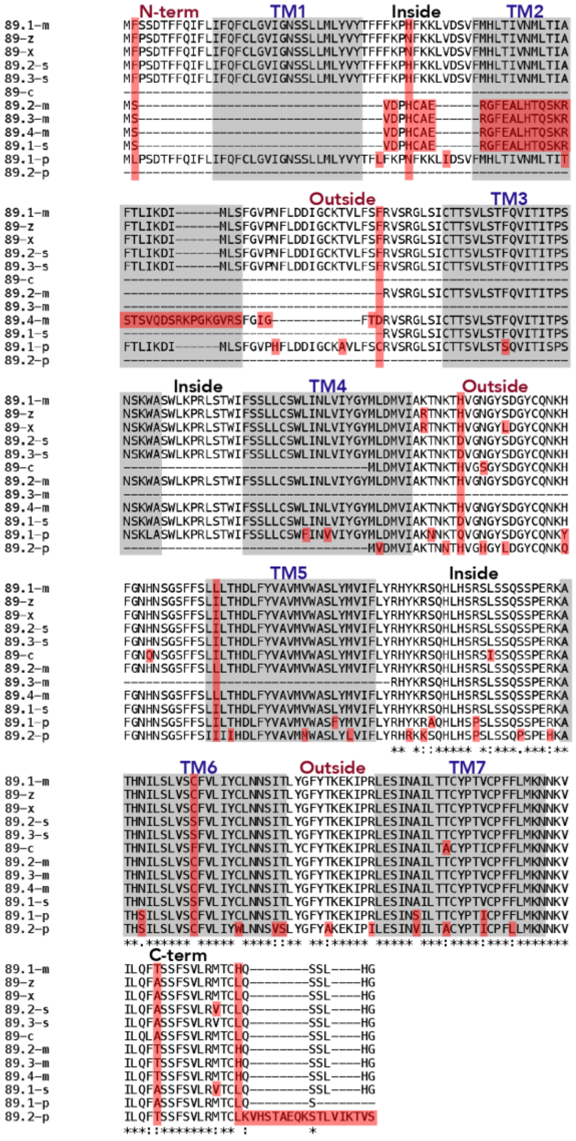


| | 89.1-m | 89.2-m | 89.3-m | 89.4-m | 89-z | 89-x | 89.1-s | 89.2-s | 89-c | 89.1-p | 89.2-p |
|---|---|---|---|---|---|---|---|---|---|---|---|
| 89.1-m | --- | 71.3 | 32.5 | 72.3 | 97.8 | 97.5 | 69.1 | 97.5 | 49.8 | 90.2 | 41.5 |
| 89.2-m | 71.3 | --- | 51.4 | **91.5** | 69.8 | 69.4 | 97.2 | 69.1 | 63.5 | 65.1 | 65.1 |
| 89.3-m | 33.1 | 51.4 | --- | 47.1 | 31.6 | 34.1 | 49.4 | 31 | 53.5 | 28.9 | 48.6 |
| 89.4-m | 74.7 | **91.5** | 47.1 | --- | 70.7 | 70.4 | **89** | 70.1 | 58.1 | 65.6 | 48.1 |
| 89-z | 98.7 | 70.4 | 32.2 | 73.8 | --- | 99.7 | 70 | 98.4 | 50.4 | 91.8 | 41.5 |
| 89-x | 98.4 | 70.1 | 35.6 | 73.5 | 99.7 | --- | 69.7 | 98.1 | 50.3 | 91.5 | 41.8 |
| 89.1-s | 69.8 | **98** | 49.8 | **89.7** | 70.6 | 70.3 | --- | 71.2 | 64.1 | 65.6 | 51.5 |
| 89.2-s | 98.1 | 69.8 | 31.9 | 73.2 | 99.4 | 99.1 | 71.2 | --- | 50.3 | 91.1 | 40.9 |
| 89-c | 51.1 | 65.1 | 55.7 | 59.6 | 51.9 | 51.6 | 65.7 | 51.6 | --- | 47.5 | 47.5 |
| 89.1-p | 93.7 | 67.9 | 30.7 | 70.4 | 94.9 | 94.6 | 68.4 | 94.6 | 49.4 | --- | 41.5 |
| 89.2-p | 45.5 | 57.3 | 54.6 | 52.6 | 45.8 | 46.1 | 56.9 | 45.2 | 80.3 | 44.8 | --- |

Identity

Similarity

**Figure S4. Alignment and pairwise comparisons of *Vmn1r89* peptide sequences**. *Mus* species indicated with letter abbrevations (*M. m. domesticus*: "**m**", *M. spicilegus*: "**z**", *M. macedonicus*: "**x**", *M. spretus*: "**s**", *M. caroli*: "**c**" and *M. pahari*: "**p**"). ***Top***: **Sequence alignment.** Identical residues indicated by asterisks. Predicted transmembrane domains shown in grey. Amino acid differences indicated in red. ***Bottom***: **Pairwise similarities and identities.** Long variants of *Vmn1r89* indicated in red. Short transcript variants among *M. m. domesticus* and *M. spretus* indicated in bold. *Vmn1r89.3-s* is not shown, as it shares 100% amino acid identity with *Vmn1r89.2-s.*

```
                N-term                 TM1                Inside      TM2
85-m     MLFPSNTILGVFVISEFCVGVLGNSLLLMLYIFNFWVKARFNRQIDPIFMHLMIVNMLTI
85-z     MLFPSNTILGVFVISEFCVGVLGNSLLLMLYIFNFWVKARFNRQIDPIFMHLMIVNMLTI
85-x     MLFPSNTILGVFVISEFCVGVLGNSLLLMLYIFNFWVKARFNRQIDPIFMHLMIVNMLTI
85.1-s   MLFPSNTILGVFVISEFCVGVLGNSLLLMLYIFNFWVKAHFNRQIDPIFMHLMIVNMLTI
85.2-s   MLFPSNTILGVFVISEFCVGVLGNSLLLMLYIFNFWVKAHFNRQIDPIFMHLMIVNMLTI
85-c     MSFPSKTLLGVIFISEFCVGIIGNSLLLVLYVYNFLVKAHFNRQIDPIFMHLMIVNMLTI
85-p     MSFPSNTLLGVFFISEFCVGVIGNSLLFVLYVYNFWVKVHFSRLIDPIFMHLMIVNILTI
         * ***:*:***:.*******::*****::**::** **.:*.* ************:***
                                 Outside                      TM3
85-m     IFAMIPYITSSFGVPSFLHDAGCKAFVYIYRVSRAMSISSTSILSTFQAITITPSNSKWA
85-z     IFAMIPYITSSFGVPSFLHDAGCKAFVYVYRVSRAMSISSTSILSTFQAITITPSNSKWA
85-x     IFAMIPYITSSFGVPSFLHDAGCKAFVYVYRVSRAMSISSTSILSTFQAITITPSNSKWV
85.1-s   IFAMIPYITSSFGVPSFLHDAGCKAFVYVYRVSRAMSISSTSILSTFQAITITPSNSKWA
85.2-s   IFAMIPYITSSFGVPSFLHDAGCKAFVYVYRVSRAMSISSTSILSTFQAITITPSNSKWA
85-c     IFSMIPYITSSFGVPSFLNSAGCKAFQYVYRVTRAMSISTTSILSSFQAITITPSNSKWA
85-p     IFAMIPYITSSFGVPSFLDNAGCKAFVYVYRVSRAMSISSTSILSTFQAITITPSNSKLA
         **:***************..****** *:***:******:*****:************ .
           Inside             TM4                       Outside
85-m     WLKPKLSKWTFCTFLFSWFINMVIYVHLIENVIAKTNYTGVNYGYSHVYCESSPVEYPNP
85-z     WLKPKLSKWTFCTFLFSWFINMVIYVHLIENVIAKTNYTGVNYGYSHVYCESSPVEYPNP
85-x     WLKPKLSKWTFCTFLFSWFINMVIYVHLIENVIAKTNYTGVNYGYSHVYCESGPVEYPNP
85.1-s   WLKPKLSKWTFCTFLFSWFINMVIYVHLIENVIAKTNYTSVNSGYSHVYCESSPVEYPNP
85.2-s   WLKPKLSKWTFCTFLFSWFINMVIYVHLIENVIAKTNYTSVNSGYSHVYCESSPVEYPNP
85-c     WLKPKLSTWTFCSFLFSWFINLVIYVQLIKTVIAKTNYTDVDYGYSHVYCGSSPVEYPNP
85-p     WLKPKLSKWTFCTFLFSWFINLVIYVHLIENVVAKTNYTDVDYGYSHVYCGSGPVEYPNP
         *******.****:********:****:**:.*:******.*: ******* *.*******
                     TM5                          Inside
85-m     GFFLGVIITRDLVLLVIMMWTSLSMVTLLYRHHKRAQHLRNPRLSSQKSPEHKATHTILL
85-z     GFFLGVIITRDLVLLVIMMWTSLSMVTLLYRHHKRAQHLRNPRLSSQKSPEHKATHTILL
85-x     GFFLGVIITRDLVLLVIMMWTSLSMVTLLYRHHKRAQHLRNPRLSSQKSPEHKATHTILL
85.1-s   GFFLGVIITRDLVLLVIMMSTSLSMVTLLYRHHKRAQHLRNQRLSSQKSPEQKATHTILL
85.2-s   GFFLGVIITRDLVLLVIMMSTSLSMVTLLYRHHKTAQHLRNQRLSSQKSPEQKATHTILL
85-c     GLFLSVIITRDLLFLVIMMWTSLYMVTLLFRHHKRAQHLRNPRLSSQKSPEHKATHTILL
85-p     GLFLSIIITRDLLFLVIMIWTSLYMVTLLYRHRKRARHLRNPSLSSQQSPEHKATRTILL
         *:**.:******::****: *** *****:**:* *:****  ****:***:***:****
               TM6        Outside             TM7          C-term
85-m     LVNCFVFFYLLNNLITLYDRFYTRGRTGNLGAIITIVASFYPTLCPFLLMNSNKMYFFPS
85-z     LVNCFVFFYLLNNLITLYDRFYTRGRTENLGAIITIVASFYPTLCPFLLMNSNKVYFFPS
85-x     LVNCFVFFYLLNNLITLYDRFYTRGRTENLGAIITIVASFYPTLCPFLLMNSNKVYFFPS
85.1-s   LVNCFVFFYLLNNLITLYDRFYTRGKTGNLGAIITIVASFYPTLCPFLLMNSNKVYFFPS
85.2-s   LVNCFVFFYLLNNLITLYDRFYTRGKTGNLGAIITIVASFYPTLCPFLLMNSNKVYFFPS
85-c     LVNCFVFFYLLNIFATLYDRFCTRERTPSLGAIIAIMASFYPTLCPFLLMNTNKI-----
85-p     LVSCFVFFYLLNNLATLYDRFYIRGKTGNLGAIITIMASFYPTLCPFLLMNNNKI-----
         **.********* : ******  * :* .*****:*:**************.**:

85-m     YIQRAIGD---------------------------------------
85-z     YIQRAIGD---------------------------------------
85-x     YIQRAIGD---------------------------------------
85.1-s   YIQRAIGD---------------------------------------
85.2-s   YIQRAIGD---------------------------------------
85-c     -ISQCISSLLVFIEPLVIDM---------------------------
85-p     -ISQCISSIHIFKESLVTDMKPRCPHQQTSLSSDLCDMLKVVWLWNPY
          *.:.*..
```

| | 85-m | 85-z | 85-x | 85.1-s | 85.2-s | 85-c | 85-p |
|---|---|---|---|---|---|---|---|
| 85-m | --- | 98.7 | 98.1 | 96.8 | 96.4 | 78.4 | 73.3 |
| 85-z | 99.4 | --- | 99.4 | 97.4 | 97.1 | 78.8 | 73.3 |
| 85-x | 98.7 | 99.4 | --- | 96.8 | 96.4 | 78.1 | 73.3 |
| 85.1-s | 97.7 | 97.7 | 97.1 | --- | 99.7 | 77.5 | 73 |
| 85.2-s | 97.4 | 97.4 | 96.8 | 99.7 | --- | 77.2 | 72.7 |
| 85-c | 85.6 | 85.6 | 85 | 84.7 | 84.4 | --- | 77.5 |
| 85-p | 79.9 | 79.6 | 79.6 | 79 | 78.7 | 83.9 | --- |

Identity

Similarity

**Figure S5. Alignment and pairwise comparisons of *Vmn1r85* peptide sequences**. *Mus* species indicated with letter abbrevations (*M. m. domesticus*: "**m**", *M. spicilegus*: "**z**", *M. macedonicus*: "**x**", *M. spretus*: "**s**", *M. caroli*: "**c**" and *M. pahari*: "**p**"). ***Top***: **Sequence alignment.** Identical residues indicated by asterisks. Predicted transmembrane domains shown in grey. Amino acid differences indicated in red. ***Bottom***: ***Vmn1r85* amino acid pairwise similarities and identities.** Conserved sequences among species closely related to the house mouse indicated in red.

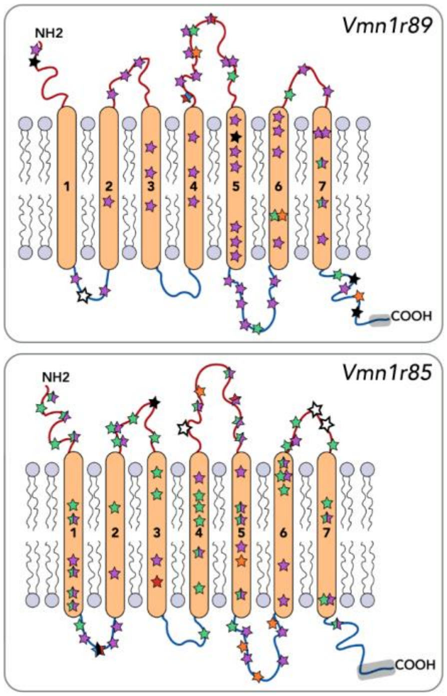


**Figure S6. Amino acid site changes in clade J/K receptors: *Vmn1r89* and *Vmn1r85***. The transmembrane protein structure of the V1Rs are depicted schematically. The location of all amino acid site changes in *Vmn1r89* and *Vmn1r85* across all 6 species are represented with stars. *Mus* species are indicated with colors and letters (*M. m. domesticus*: black; *M. spicilegus*: blue; *M. macedonics*: red; *M. spretus*: orange; *M. caroli*: green; *M. pahari*: purple). Stars with multiple colors indicate an amino acid change present in multiple species at that site. White stars indicate *highly variable sites* (several variable amino acid changes are present across multiple species). The percentage of amino acid site differences within each region (transmembrane, inside, or outside the membrane) across all species are indicated in table below. Short *Vmn1r89* transcripts were excluded from this analysis.

**% Amino Acid Differences by Protein Region**

| | Transmembrane | Inside (C-terminus & intracellular loops) | Outside (N-terminus & extracellular loops) |
|---|---|---|---|
| ***Vmn1r89*** | 15.3 | 20.0 | 22.9 |
| ***Vmn1r85*** | 22.7 | 20.6 | 29.4 |
